# Performance anxiety is associated with context-dependent reward and punishment learning biases in skilled performers

**DOI:** 10.1101/2025.03.31.646422

**Authors:** Andrea Erazo Hidalgo, Lisa Pearson, Takanori Oku, Yudai Kimoto, Shinichi Furuya, María Herrojo Ruiz

**Author notes:** **Corresponding author:** Maria Herrojo Ruiz,. Address: School of Mind, Body and Society, Goldsmiths, University of London. Lewisham Way, New Cross, London SE14 6NW (UK). AEH and MHR contributed equally to this work. **Author contributions:** Conceptualization: MHR, LP, SF. Methodology: MHR, AEH, TO, YH, LP. Formal analysis: MHR, AEH. Investigation: AEH, YH, MHR. Data curation: AEH, MHR. Writing original draft/visualization: AEH, MHR. Review and editing: all authors. Supervision: MHR, SF. Funding: MHR, AEH, SF.

## Abstract

Performance anxiety (PA) is common in high-stakes settings, from public speaking to the performing arts, yet its mechanisms remain poorly understood. Using behavioural analysis, computational modelling, and electroencephalography, we tested whether PA biases learning from reward and punishment under varying task uncertainty. Across three experiments with 95 skilled pianists, participants learned hidden melody dynamics through graded reward or punishment feedback. Bayesian hierarchical regression showed that increasing PA levels were associated with faster reward learning under high uncertainty (large action space) but faster punishment learning under low uncertainty (reduced action space). These learning biases—reflecting the interaction between PA and reinforcement type—were associated with reinforcement-driven modulation of motor variability and frontal theta (4–7 Hz) activity encoding feedback changes and signalling motor adjustments. The findings reveal that predisposition to PA in skilled individuals modulates reward- and punishment-based learning in a context-dependent manner governed by task uncertainty.

## Introduction

Performing in high-stakes, socially evaluative settings—where individuals are judged on their abilities—is a fundamental challenge in human behaviour, spanning domains as diverse as sports, public speaking, and the performing arts. While some individuals thrive in those settings, others experience performance anxiety (PA)—a debilitating condition characterised by anxious apprehension towards performance^1^. PA affects between 25% and 40% of professionals and students across domains^2–5^, significantly impacting health and career trajectories, yet its underlying neurocognitive mechanisms remain poorly understood.

PA is characterised by altered physiological, cognitive, and affective states^3,4,6–9^, often impairing performance in critical moments. Competitions and stage performances heighten state anxiety, disrupting cardiorespiratory rhythms and motor control^6–8,10^. Laboratory studies show that PA increases muscle stiffness, impairs memory retrieval, and disrupts the automatic execution of well-learned actions^11–13^. Despite these advances, a major gap remains in understanding how PA interacts with fundamental learning processes.

Anxiety disorders are increasingly conceptualised as disorders of learning and decision-making, particularly under uncertainty^14–16^, where information is incomplete or the environment is unstable. A prevailing hypothesis is that anxiety is associated with negative learning biases^16–18^, whereby individuals exhibit a greater reliance on negative outcomes to update their behaviour or beliefs. Computational studies have shown that clinically anxious individuals learn faster from punishment than from reward^17,18^, an effect attributed to biased attention for threats and suppression of reward-seeking behaviour^16,19,20^. Variations in these patterns associate cognitive symptoms of anxiety with faster threat learning, while physiological symptoms increase safety learning^21^. Decision-making studies further show that abnormal learning processes in anxiety intensify as uncertainty increases^22–26^. Similarly, in motor learning tasks involving large, intrinsically uncertain continuous action spaces, state anxiety attenuates reward learning^27^. Based on these findings, we hypothesise that PA promotes negative learning biases, leading affected individuals to rely more on punishment than reward to guide adaptation during performance—an effect that may be exacerbated under uncertainty.

In the motor domain, reward and punishment differentially modulate learning. While punishment increases learning rates during sensorimotor adaptation^28,29^, reward improves retention of adaptation and motor skills^28,30,31^. However, inconsistencies in these findings indicate task-dependent effects^32,33^. Faster punishment learning has been explained by greater trial-by-trial motor variability and larger motor updates following negative outcomes^28,31^. After unsuccessful actions, increased task-related variability promotes exploration, enabling the sensorimotor system to more rapidly identify successful actions^34–36^.

Motor variability arises from several sources^37,38^, including neuromotor and planning noise, along with exploratory variability, which is particularly sensitive to outcomes about success and failure. By dissociating reinforcement effects from behavioural autocorrelations, recent studies demonstrate that poor outcomes causally increase exploratory variability to improve learning^36,39^. Computational modelling complements these findings by revealing how agents adjust different sources of motor variability^36,39^, balancing exploratory variability against motor noise. Together, these approaches provide a framework to test our second hypothesis: that, in the motor domain, PA biases learning from reward and punishment through altered regulation of motor variability.

To identify the neural mechanisms underlying the hypothesised learning biases and altered motor variability regulation in PA, we examined electroencephalography (EEG) oscillations. Prefrontal and sensorimotor beta oscillations (13–30 Hz) are key modulators of motor learning^40–43^, including reward-based learning^27,44,45^, with beta attenuation post-feedback contributing to updating motor plans^42,45,46^. In line with this, increased beta activity in state-anxious individuals has been linked to attenuated reward processing, impairing the updating of motor predictions^27^. Additionally, frontal midline theta oscillations (4–7 Hz) have been implicated in adaptive control, adjusting behaviour under uncertainty by facilitating switching and exploration in reinforcement learning^47–50^. Theta is also associated with a predisposition to anxiety and heightened responses to punishment^51^, suggesting a role in mediating reinforcement-learning biases in PA.

Beta and theta oscillations in these settings have been explained by activations in the anterior cingulate cortex (ACC), prefrontal cortex (PFC), hippocampus, and striatum^52,53^—regions crucial to decision-making, learning under uncertainty, and anxiety^14,54–57^. The striatum, a key structure in reinforcement-based motor learning^58^, is part of the cortico-basal ganglia-thalamo-cortical circuits, which are proposed to regulate motor variability^35,37,39,59^. In the cortex, the PFC tracks hidden task states by predicting observations during reward-guided decision-making^60^, complementing the role of the basal ganglia in learning via reward prediction errors. Here, we tested whether beta and theta modulation reflects changes in reinforcement-based motor learning in PA. We predicted a more pronounced beta attenuation during punishment learning, associated with faster avoidance learning, and enhanced medial frontal theta, encoding control signals for greater behavioural adjustments and increased motor variability regulation under punishment. If confirmed, these EEG dynamics would mark a neurophysiological signature of learning biases in PA, reflecting maladaptive learning mechanisms that may undermine skilled performance.

Despite extensive evidence linking anxiety to learning biases, investigating these mechanisms in highly trained individuals with a predisposition to PA has remained challenging. This shortfall is partly due to methodological constraints in assessing skilled performance, which requires simultaneous recording of rich performance data and neural activity. To address this, our study focused on skilled pianists, enabling a quantitative assessment of reinforcement learning and motor variability in expert sensorimotor performance.

Across three experiments with 95 pianists, we examined how trait PA influences learning from reward and punishment and reinforcement-driven motor variability, with neural correlates assessed in Experiment 1. We used performance learning tasks requiring pianists to adapt keystroke dynamics (intensity or loudness) to uncover hidden target dynamics in melodies under graded reward or punishment reinforcement. Contrary to our first hypothesis, Experiments 1 and 2 revealed that increasing PA levels were associated with faster reward learning, whereas lower PA levels corresponded to greater reliance on punishment feedback. In Experiment 3, which involved a reduced action space reflecting lower task uncertainty, these learning biases reversed. Across experiments, these learning biases were associated with reinforcement-driven regulation of motor variability. At the neural level, theta activity parametrically encoded unsigned differences in graded reinforcement feedback and upcoming motor adjustments, accounting for the observed learning biases.

These findings indicate that predisposition to PA manifests as biases in learning from reward and punishment, accompanied by altered regulation of reinforcement-driven motor variability and associated with changes in frequency-domain activity. The reversal of learning biases across tasks with large versus reduced action spaces—reflecting high versus low uncertainty—highlights their context-dependent nature and underscores the role of uncertainty in PA.

## Results

To evaluate the dissociable effects of reward and punishment on learning in skilled performers, we developed a performance learning task adapted from previous reward-based motor learning research^27^ and used it to collect behavioural and EEG data from a cohort of highly trained pianists (N = 41). The data are available online (see **Data Availability Statement**).

Participants played two piano melodies designed for the right hand on a digital piano (**Figure 1A**). The task entailed varying the dynamics (the pattern of keystroke velocity or loudness) with the aim of uncovering the melody’s specific hidden target dynamics. Participants were informed that the target dynamics deviated from the natural flow of the melodies (**Figure S1**) and would not correspond with their initial expectations, requiring exploration to uncover the solution (**Methods**).

**Figure 1.**
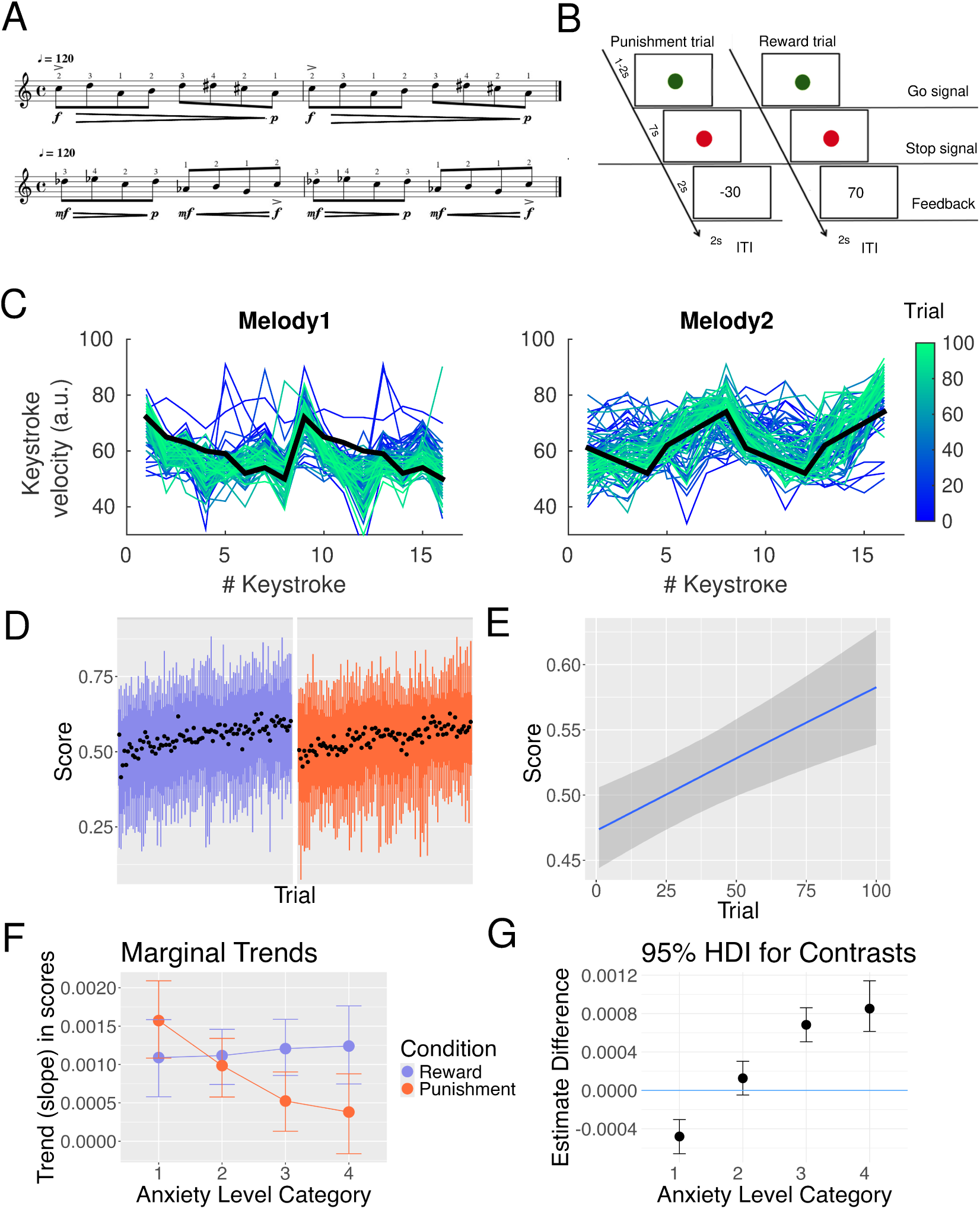
Task and Performance Analysis for Experiment 1. **A.** Participants (N = 41 skilled pianists) played two right-hand piano melodies on a digital keyboard, adjusting their keystroke dynamics (intensity of key press) to uncover a hidden target dynamics pattern. Beneath the musical scores, the flow of the target dynamics is represented in musical notation, with *p* denoting *piano*, *f forte*, and *mf mezzo-forte*. **B.** Trial timeline. Each trial yielded graded reinforcement feedback in the form of reward (0–100) or punishment (– 100–0) scores, separately in each reinforcement condition. Feedback was a function of the proximity of performed dynamics to the hidden target. **C.** Example of melody dynamics performed by one participant (graded blue to green for trials 1 to 100), with target dynamics denoted by the bold black line. **D.** Score progression across trials in the group, representing mean with 66% and 95% confidence intervals for the reward (purple) and punishment (orange) conditions. **E.** Bayesian multilevel beta regression modelling revealed a credible positive effect of trial progression on score (posterior median slope = 0.00441; 95% credible interval [0.00243, 0.00651]; log-odds scale), reflecting learning to approach the target dynamics. **F–G.** Marginal trends. A three-way interaction between reinforcement condition, trial, and trait levels of performance anxiety (PA) showed a credible dissociation: lower-PA participants learned faster under punishment than reward (reward minus punishment median slope estimate: -4.81 × 10^-4^, 95% highest density interval, HDI [-6.60, -3.04] × 10^-4^). By contrast, medium-high and higher-PA individuals showed steeper learning under reward (median slope difference: 6.83 [5.06, 8.61] × 10^-4^ and 8.52 [6.14, 11.42] × 10^-4^, respectively). Coloured circles denote median estimates, and shaded intervals indicate 95% HDI of the posterior distribution of median trends for reward and punishment conditions, as well as for the contrast in panel G (reward minus punishment difference estimates).

After each trial, participants received graded reinforcement feedback, presented in separate reward (scores 0–100) and punishment (–100 to 0) condition blocks of 100 trials each ( **Figure 1B**). This feedback reflected their overall proximity to the target dynamics pattern, calculated as a single summary score comparing the full vector of performed keystroke velocities to the target dynamics vector for that melody (**Figure 1C; Methods**). The goal was to infer the hidden dynamics solution and maximise the average score across trials—coupled to a monetary incentive, by either increasing gains in the reward condition or minimising losses in the punishment condition. Concurrently, EEG and MIDI (Musical Instrument Digital Interface) performance data were recorded.

### Bayesian workflow of performance analysis

To assess the effects of reinforcement condition on learning and its interaction with trait performance anxiety (PA), we used Bayesian multilevel regression modelling^61^. PA was evaluated using the validated Kenny music performance anxiety (MPA) Inventory^62^. See simulations for sample size estimates in **Figures S2-S3**, which recommended a sample of 40 participants.

Following the principled Bayesian workflow^61^, we constructed Bayesian beta regression models of feedback scores (rescaled to 0–1) over trials, analysing the effects of reinforcement, PA levels, and their interaction (**Methods**; **Table S1).** Beta regressions were parametrised by the mean *μ* and precision *ϕ* of the score distribution^63^. Prior predictive checks with simulated data confirmed that model behaviour aligned with domain expertise (**Figure S4**).

The best-fit model included interactions between reinforcement condition, a monotonic function of PA categorical levels^64^, and trial progression, along with random intercepts and slopes for subjects (model M6, **Table S2;** Leave-one-out cross-validation^65^, LOO-CV). This model demonstrated good convergence and robust predictive accuracy (**Figure S5**, **Supplementary Materials**).

Scores increased across trials, with a positive effect of 0.00441 per trial on the log-odds scale (95% credible interval, CrI: [0.00243, 0.00651]), equivalent to an increase of 10 points over 100 trials. This validates that participants progressively approached the target dynamics (**Figure 1DE; Table S2)**. Score consistency also increased (greater precision parameter *ϕ*), and a credible three-way interaction between PA, condition and trial on the scores was observed.

Additional analyses of marginal trends (slopes) revealed that, overall, reward led to steeper learning slopes than punishment (reward – punishment: 2.97 × 10^-4^, 95% HDI [1.78, 4.18] × 10^-4^). Further analysis of the interaction showed distinct credible PA-dependent effects of reinforcement condition on the median trend of scores across trials (**Figure 1FG**; effects on the percentage point scale). Low-PA participants learned faster to avoid punishment (negative median slope difference, reward – punishment: -4.81 × 10^-4^, 95% highest density interval, HDI [-6.60, -3.04] × 10^-4^). Conversely, individuals with medium-high to high PA learned faster to maximise reward (median slope difference: 6.83 [5.06, 8.61] × 10^-4^ and 8.52 [6.14, 11.42] × 10^-4^, respectively), with the most pronounced difference at the highest PA level.

These results demonstrate that although learning rates were generally higher under reward, they were distinctly modulated by reinforcement condition as a function of PA, exhibiting a monotonic shift from faster punishment learning at low PA to faster reward learning at high PA. The interaction effects on learning trends did not extend to median scores (**Figure S6**). These findings were replicated in a second experiment with an independent sample of 18 highly trained pianists, confirming similar interaction effects on learning slopes (**Figure 2**; **Figure S7, Table S3)**. As in Experiment 1, lower-PA participants learned faster under punishment than reward, whereas medium- to high-PA individuals showed progressively steeper learning under reward. Overall, reward learning was also faster than punishment learning.

**Figure 2.**
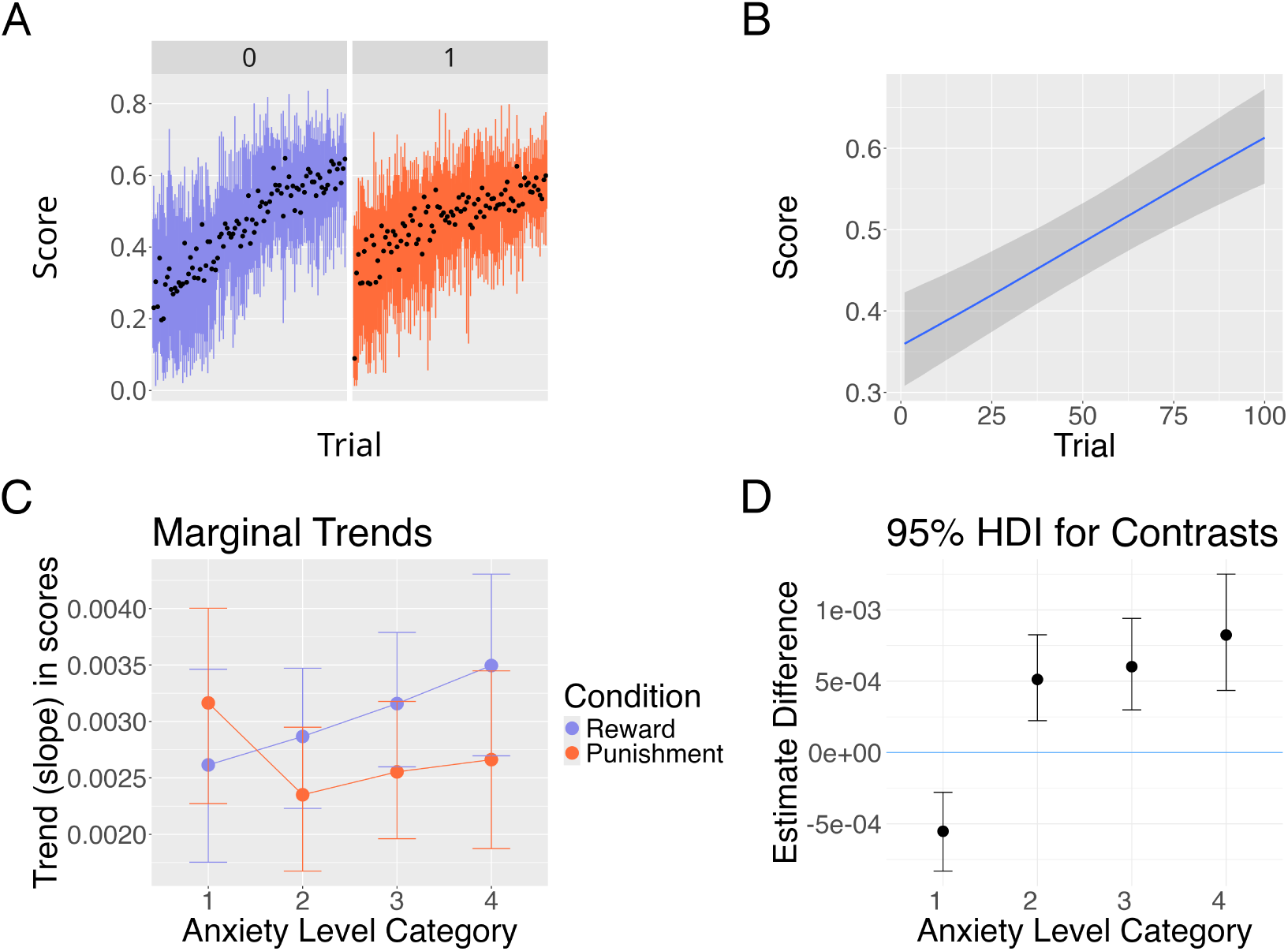
Marginal effects in Experiment 2 (N = 18). A–B. Marginal trends. As in Experiment 1, a three-way interaction between reinforcement condition, trial, and trait PA levels showed a credible dissociation: lower-PA participants learned faster under punishment than reward (reward minus punishment median slope estimate: -5.52 × 10^-4^, 95% highest density interval, HDI [-8.32 -2.79] × 10^-4^). By contrast, medium, medium-high and higher-PA individuals showed steeper learning under reward (median slope difference: 5.12 [2.23 8.25] × 10^-4^, 6.02 [2.99 9.40] × 10^-4^ and 8.24 [4.35 12.50] × 10^-4^, respectively). Coloured circles denote median estimates, and shaded intervals indicate 95% HDI of the posterior distribution of median trends for reward and punishment conditions, as well as for the contrast in panel B (reward minus punishment difference estimates)

Given previous findings that cognitive (worry) and somatic (physiological) trait anxiety symptoms can influence learning biases differently^21^, we conducted control analyses to determine if distinct PA components differentially influence learning biases. Bayesian multilevel modelling revealed that the model incorporating somatic PA subscores^66,67^ provided a better fit than the model including cognitive PA (negative cognitions), replicating the interaction effects (Experiments 1 and 2: **Figures S8-S9, Table S4**). This suggests that learning biases in skilled performers are better explained by the debilitating physiological dimension of PA.

### Learning biases in performance anxiety are associated with changes in motor variability

To assess our second hypothesis, we examined trial-by-trial changes in motor variability. Our task involved a large continuous action space, requiring participants to infer hidden melody dynamics across 16 (8 × 2) keystroke velocities using reinforcement feedback. In such environments, learning can be effectively guided by reinforcement-driven regulation of motor variability^35–37,39^: variability increases following unsuccessful outcomes to promote exploration, and decreases after successful outcomes to stabilise performance.

To assess variability in keystroke velocity, we first transformed the trial-wise 16-dimensional velocity vector into a scalar variable, *ΔV^n^*, representing the magnitude of change in velocity patterns between consecutive trials (*n*–1 and *n*). *ΔV^n^* was computed as the normalised sum of absolute differences across corresponding keypresses *k* between the two trials (eq.[1] in **Methods**; **Figure 1C**)^68^. Following ref.^36^, variability was then assessed by calculating the variance of *ΔV^n^* values within moving windows of five trials.

As expected^35,36,68^, participants increased variability more following poor than high outcomes (median split of scores; **Figure 3A-B**). Slow fluctuation trends, potentially reflecting autocorrelations in performance^40,69–73^, were also evident in the variability function in renditions preceding and following conditioned trials^36^ (relatively low or high scores; **Figure 3B**).

**Figure 3.**
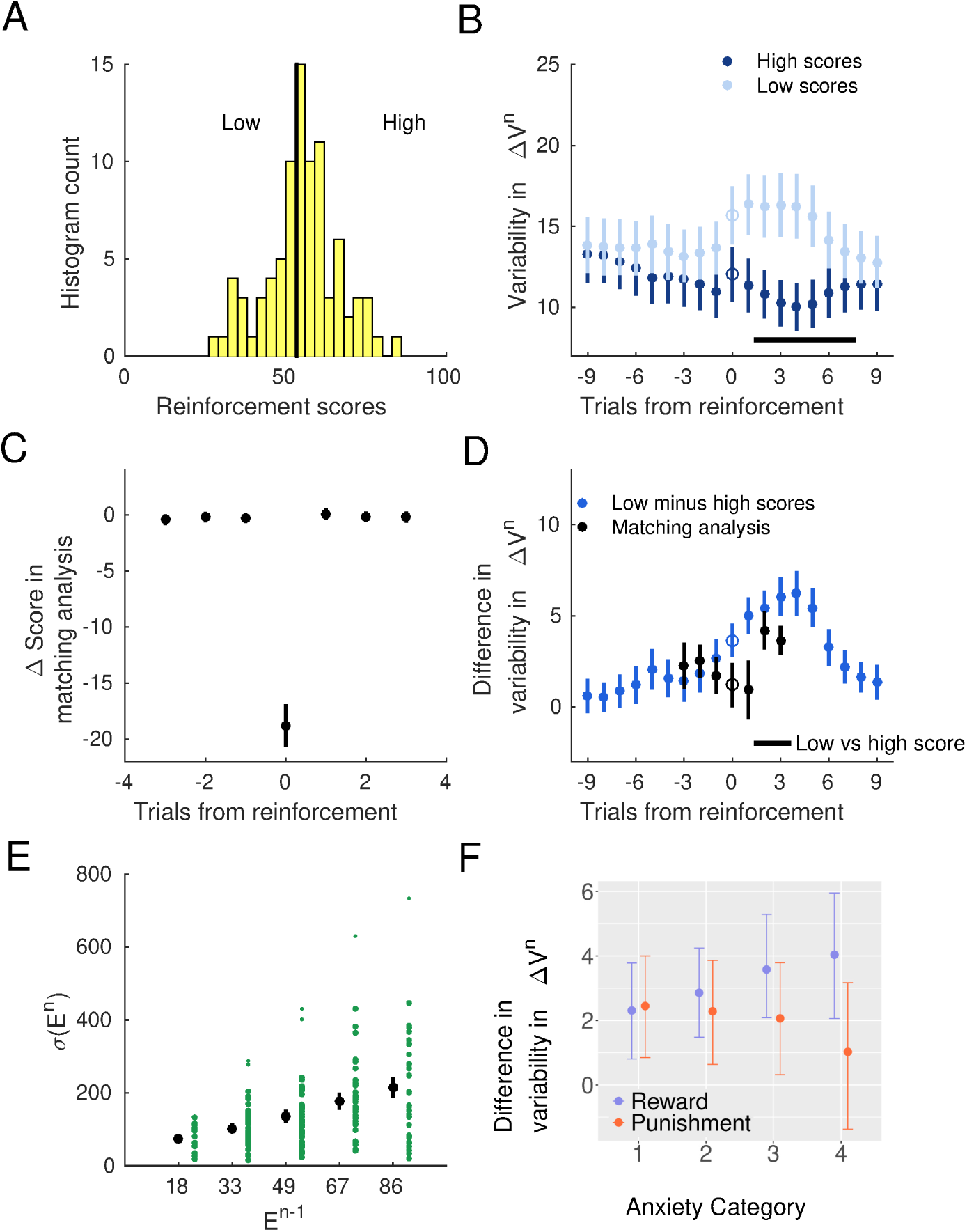
Reinforcement-related modulation of motor variability in Experiment 1. **A.** Histogram of reinforcement scores (in one example participant and condition) illustrating the median split used to define low and high score conditions. **B.** Time course of task-relevant motor variability surrounding high (dark blue) and low (light blue) score trials (relative to median split), aligned to the conditioned trial at position 0. Motor variability was assessed using the variance of *ΔV^n^*values across running windows of five trials, where *ΔV^n^* represents the normalised sum of absolute differences in keystroke velocity at each of the 16 positions in a melody between consecutive trials (*n–1* to *n*). Variability was significantly higher following low scores compared to high scores at positions +1 to +3 (N = 41; paired permutation test; *P_FDR_*= 0.001, significant effects denoted by the horizontal bar at the bottom). Large coloured dots indicate means, with error bars denoting ± SEM. **C.** Statistical matching analysis: we selected trials of low and high scores (median split) that were preceded and followed by similar reinforcement values (and thus performance). The black dots represent the mean performance score difference (SEM) at each trial. **D.** Using trials obtained from the matching analysis (black), we found that motor variability was significantly greater at lags +1 to +3 following conditioned trials (position 0) associated with low compared to high scores (*P_FDR_* = 0.0038; dots represent mean, and bars SEM). The difference in uncorrected (all trials) motor variability between low and high scores, not accounting for performance autocorrelations, as shown in panel B, is depicted in blue. **E.** Larger deviations from target velocity patterns—measured as the norm of vector differences and represented as unsigned error E^n-1^—were followed by greater subsequent reinforcement-related variability, *σ(*E^n^). In our task, larger deviations were associated with lower scores. **F.** The difference in motor variability following poor versus good outcomes, labelled *VarDiff* in the main text and obtained from the matching analysis trials, was modulated by the interaction between PA categorical level and reinforcement condition. A negative estimate (-1.06, 95% CrI: [-2.13, -0.01]) indicated that punishment, compared to reward, reduced variability following poor outcomes as PA levels increased. Coloured dots represent posterior point estimates (reward in purple, and punishment in orange) and bars denote 95% CrI.

To dissociate performance autocorrelations from reinforcement-dependent variability and determine whether trial outcomes influence motor variability in our task, we implemented two validated approaches: statistical matching analysis and computational generative modelling^36,39^ (**Methods**, and next section). Statistical matching analysis indicated that poor outcomes led to a gradual increase in variability over 3-4 trials (**Figure 3D**). This increase was significantly greater than that following high scores (paired permutation test; *P_FDR_* = 0.0038; non-parametric effect size estimator, Δ_dep_ = 0.74, CI = [0.65, 0.89]). Separately, we observed that larger deviations from target velocity patterns (lower scores) resulted in greater subsequent reinforcement-related variability (**Figure 3E**), in line with previous work^39^.

To determine whether the observed learning biases (**Figure 1C-D**) were associated with changes in the regulation of motor variability, we applied Bayesian Gaussian linear modelling to analyse *VarDiff*—the difference in variability following poor minus good outcomes—as a function of PA category, reinforcement condition, and their interaction. The model demonstrated good convergence (**Table S5**) and revealed a credible interaction between both variables. A negative estimate (-1.06, 95% CrI: [-2.13, -0.01]) indicated that punishment, compared to reward, reduced variability following poor outcomes as PA levels increased (**Figure 3F**). This effect was most pronounced for high PA individuals, where poor outcomes did not increase variability under punishment (95% CrI overlapping zero). No other effects were observed (**Table S5**).

Thus, controlling for behavioural autocorrelations, our analysis provided consistent evidence that reinforcement-driven motor variability is associated with learning biases in skilled performers as a function of PA. Moreover, high-PA individuals exhibited the most contrasting responses to reward and punishment, with preserved motor variability regulation under reward but blunted responses under punishment.

Control analyses defining low and high scores based on relative trial-to-trial score changes confirmed that motor variability regulation was primarily driven by poor outcomes below the median of the score distribution (**Supplementary Materials**). Notably, these low scores were distributed across the entire session rather than concentrated in earlier trials (**Supplementary Materials**), ruling out confounds from early-session effects.

#### Generative model dissociating reinforcement-sensitive and autocorrelated behaviour in motor variability

The previous results suggest that keystroke dynamics were influenced by reinforcement-driven changes in motor variability, alongside slower autocorrelations in performance. Similar patterns have been observed in humans and non-human primates^39^, as well as rodents^36^. In such settings, a control strategy involves counteracting autocorrelations and baseline motor noise—which both reduce reward rates—by increasing variability to explore and identify reinforced solutions^34,38,39,74,75^.

To investigate whether this control strategy underlies the learning biases associated with PA, we employed a reinforcement-sensitive Gaussian process^39^ (RSGP; **Methods**). The RSGP models behavioural time series by incorporating long-term autocorrelations and short-term reinforcement effects on motor variability via two kernels (**Figure 4A**). Each kernel is defined by two hyperparameters: a characteristic length-scale (*l*), indicating the length of trial-to-trial dependencies, and an output scale (*σ*^2^), quantifying the magnitude of variability attributed to that process. In this framework, *σ*_RS_^2^ reflects the *latent* contribution of short-term, reinforcement-sensitive processes to motor variability, while *σ*_SE_^2^ (squared exponential kernel) reflects variability due to longer-term autocorrelations. An additional term, *σ_0_*^2^, accounts for unstructured, time-independent baseline noise, which may reflect each individual’s intrinsic motor noise.

**Figure 4.**
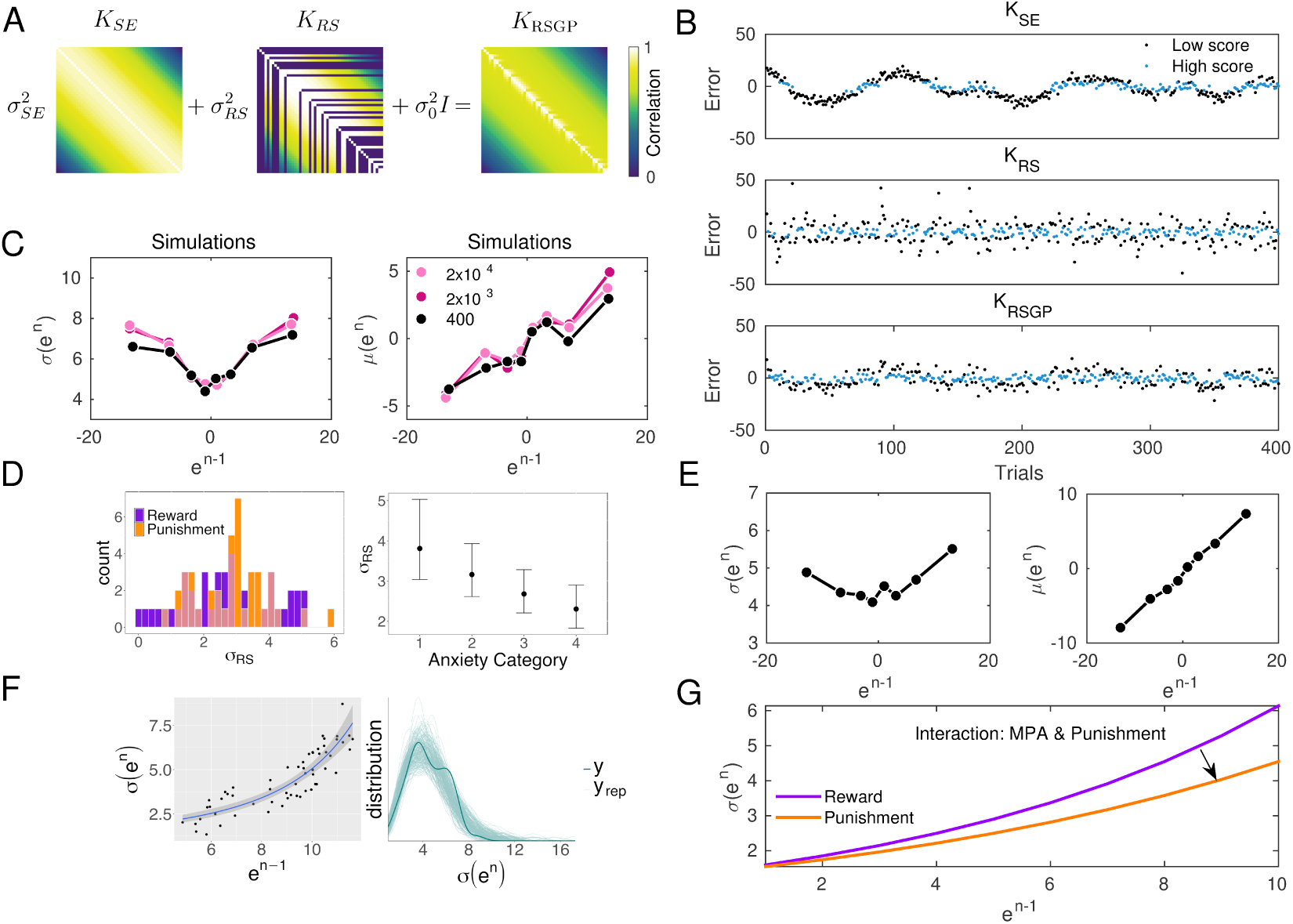
Reinforcement-sensitive Gaussian process dissociates the effects of autocorrelations and the short-term influences of reinforcement on motor variability in Experiment 1. **A.** Correlation structures of the squared exponential (K_SE_), reinforcement-sensitive (K_RS_), and combined reinforcement-sensitive Gaussian process (K_RSGP_) kernels used to model the time series of deviations between the produced and target keystroke velocity vectors, the trial-wise signed error *e^n^*. **B.** Simulated trial-wise errors (*e^n^*) under K_SE_, K_RS_, and the combined K_RSGP_. Black dots indicate trials associated with low scores (median split), while blue dots correspond to high-score trials. **C.** Simulations from the generative RSGP model reproduce key empirical patterns from previous work^39^. Left: standard deviation of error (*σ(e^n^)*) shows a U-shaped dependence on the error on the previous trial (*e^n-1^*). Right: mean error (*μ(e^n^)*) increases linearly with *e^n-1^*. Coloured lines (dots) reflect the number of samples used in the simulations. **D.** Left: distribution of *σ*_RS_ estimates across participants, split by reinforcement condition (orange for punishment, purple for reward). Right: Bayesian regression (log-normal family) revealed that *σ*_RS_ declined with increasing performance anxiety (PA) category (posterior estimate in log scale: -0.17, 95% CrI = [-0.29, -0.06]). This implies that the latent contribution of reinforcement-sensitive variability to *e*^n^ decreased as PA increased. **E.** RSGP fit to empirical data replicates simulation results from C: *σ(e^n^)* (left) and *μ(e^n^)* (right) as functions of *e^n-1^*. Mean and SEM are shown; however, SEM values are very small and not visually noticeable. **F.** Left: Exponential fit (line, 95% CrI shaded) to the relationship between *σ(e^n^)* and |*e^n-1^*|. Right: posterior predictive distribution of *σ(e^n^)*, with individual posterior draws (light green) and empirical data (dark green). **G.** Illustration of the interaction between PA category and reinforcement condition on parameter *b*₂ in the exponential relationship between *σ(e^n^)* and |*e^n-1^*|. Posterior estimate: *b_2_* (-0.03, 95% CrI = [-0.07, -0.01]). This revealed that the exponential growth in observed motor variability was attenuated under punishment (orange) relative to reward (purple) as PA levels increased (arrow), suggesting reduced sensitivity to prior error under punishment.

In line with ref.^39^, we used the trial-wise *signed* error (*e^n^*)—the difference between the produced and target keystroke velocity vectors—as the variable predicted by the RSGP at trial *n* (**Figure 4B**; **Methods**; Eq. 2), assuming a zero mean Gaussian process. *Observed total* motor variability was quantified as the standard deviation of the error distribution, *σ(e^n^),* and examined as a function of the error on the previous trial, *e^n-1^* .

Simulations confirmed reliable recovery of model parameters (*l*_SE_, *l*_RS_, *σ*_SE_, *σ*_RS_, *σ*_0_; **Table S6)** and generated predictive distributions of ***e*^n^** per trial, based on reinforcement history and prior error *e*^n-1^, characterised by the mean *μ(e^n^)* and standard deviation *σ(e^n^)*. The simulations revealed a U-shaped relationship between *e^n-1^* and *σ(e^n^)*, while *μ(e^n^)* increased linearly with *e^n-1^*, consistent with previous findings^39^ (**Figure 4C)**.

Fitting the RSGP to empirical data (separately for each condition) revealed that the autocorrelation kernel had a significantly longer length-scale (*l*_SE_ = 10.12 [0.8]) and larger output scale (*σ*_SE_ = 3.80 [0.2]) than the reinforcement-sensitive kernel (*l*_RS_ = 2.83 [0.3]; *σ*_RS_ = 2.72 [0.1]; *P_FDR_* = 0.0002; Δ_dep_ = 0.80, CI = [0.73, 0.88] for *l*, Δ_dep_ = 0.76, CI = [0.60, 0.79] for *σ*). Thus, autocorrelation effects spanned ∼10 trials, while reinforcement effects decayed after ∼3 trials. The output scale of the independent baseline noise was *σ*_0_ = 2.142 [0.08]).

Bayesian regression modelling (log-normal family) demonstrated a credible negative effect of PA category on *σ*_RS_ (log scale: -0.17, 95% CrI = [-0.29, -0.06]), indicating that the latent contribution of reinforcement-sensitive variability to *e*^n^ decreased with increasing PA, regardless of reinforcement type. No credible effects were observed for *σ*_SE_ (**Supplementary Materials; Figure 4D**). Interestingly, however, *σ*_0_ decreased with increasing PA, suggesting that greater PA was associated with reduced baseline noise (log scale: -0.14, 95% CrI = [-0.24, -0.04]).

Simulating ***e*^n^** from individual parameter estimates replicated the empirical U-shaped relationship between *σ(e^n^)* and *e^n-1^* and the linear increase in *μ(e^n^)* with *e^n-1^* (**Figure 4E**). This supports that total motor variability increased more after trials with lower scores (greater *e^n-1^*). To link these results to **Figure 3E**, we transformed *e^n-1^* into positive values,|*e^n-1^|,* and modelled the nonlinear relationship between *σ(e^n^)* and |*e^n-1^|*. An exponential Bayesian regression model (*σ(e^n^)* = *b_1_* exp(*b_2_* |*e^n-1^|*)) best explained the data (LOO-CV; **Figure 4F**; **Supplementary Materials**), with non-zero posterior estimates for both *b*_1_ and *b*_2_. A credible interaction between PA category and reinforcement condition on *b_2_* (-0.03, 95% CrI = [-0.07, -0.01]) indicated that increasing PA levels dampened the exponential growth of *σ(e^n^)* under punishment compared to reward (**Figure 4G**), suggesting progressively reduced sensitivity of observed motor variability to prior error under punishment with higher PA.

### Electroencephalography markers underlie learning biases and motor variability regulation

Having identified that increasing PA levels in skilled pianists were associated with greater motor variability following poor outcomes under reward, but a progressively blunted modulation under punishment, we next examined the neural processes underlying these behavioural effects across the theta and beta bands. We assessed frequency-domain amplitude changes related to processing graded feedback and regulating motor variability in keystroke dynamics using validated linear convolution models for oscillatory responses^76^. The frequency-domain general linear model (GLM) included parametric regressors for trial-wise unsigned changes in feedback scores and the scalar variable *ΔV^n^* denoting changes in keystroke dynamics from the current to the next trial^68^. A discrete regressor was included for feedback onset (see **Methods**). Alternative GLM models using different score representations (graded scores 0–1, signed score differences) were discarded due to regressor collinearity, which risked model misspecification (**Methods**).

Theta-band activity significantly increased more following punishment than reward feedback (**Figure 5A**; P_FWER_ = 0.021, cluster-based permutation test; **Figure 5B**; N = 39), consistent with previous work^47,49^. This effect emerged between 0.2–0.45 s in frontocentral electrodes. Beyond this feedback-related response, theta activity parametrically tracked unsigned score changes but in opposite directions for reward and punishment: it increased with greater score changes under reward but decreased under punishment, with a significant between-condition difference (P_FWER_ = 0.010; 0.2–1 s; **Figure 5C**). This effect was observed in left frontocentral and right centroparietal electrodes (**Figure 5D**). Additionally, theta amplitude reflected upcoming motor adjustments in a reinforcement-dependent manner: it increased with greater dynamics changes *ΔV^n^* in the next trial under reward but decreased under punishment, with a significant difference between conditions (P_FWER_ = 0.009; 0.2–0.9 s; **Figure 5E**), spanning midline frontal and left central electrodes (**Figure 5F**).

**Figure 5.**
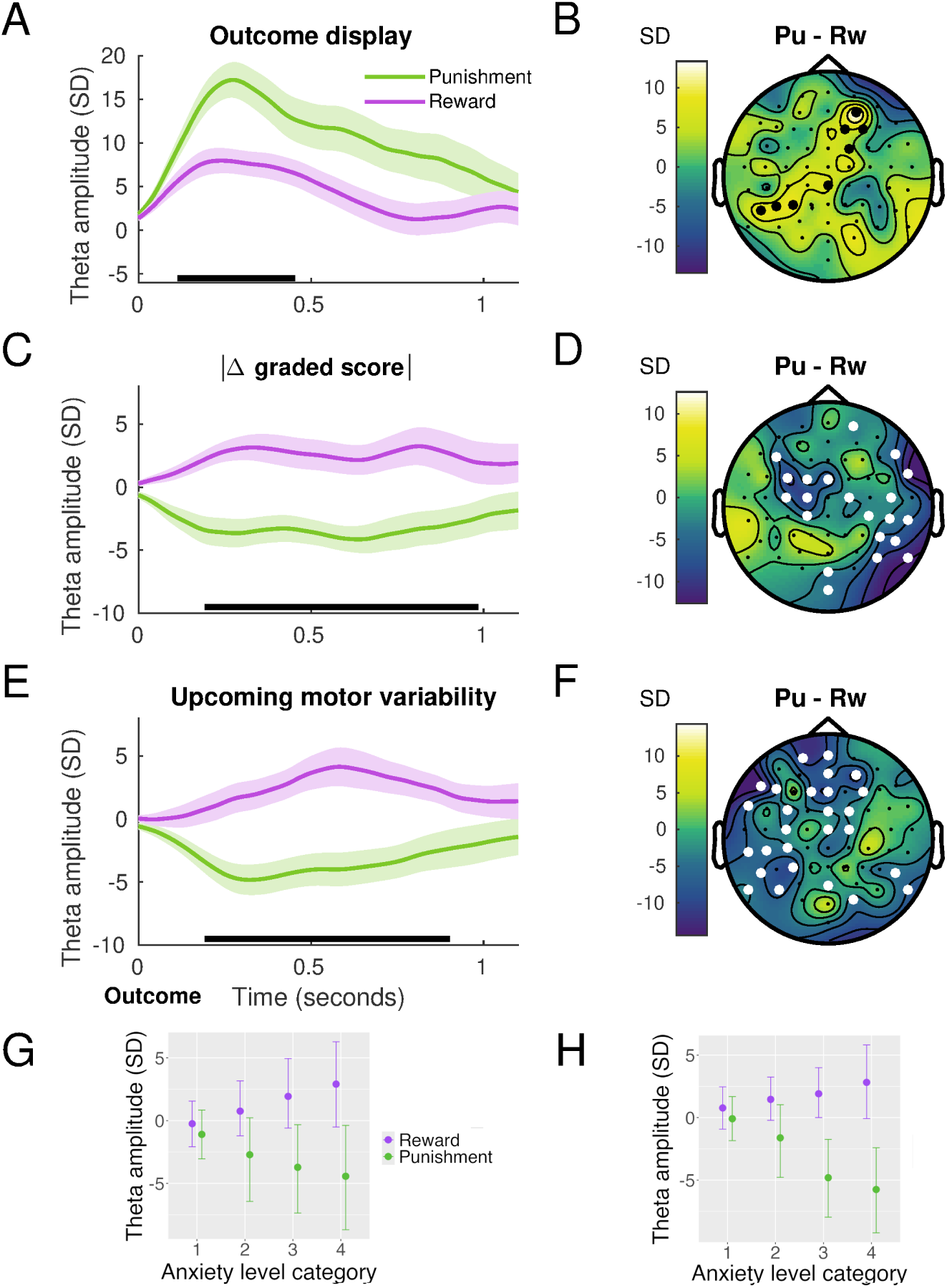
Theta activity is modulated by unsigned score differences and the amount of upcoming motor adjustments in a reinforcement-dependent manner. **A.** Time course of feedback-locked theta amplitude (4–7 Hz) in response to outcome onset (reward, magenta; punishment, green), showing greater theta power following punishment than reward between 0.2–0.45 s. Shaded regions denote ±1 SEM; black horizontal bars indicate significant cluster intervals (N = 39; cluster-based permutation test, P_FWER_ = 0.021, FWER-corrected). **B.** Topographic map of the difference (punishment – reward) in theta amplitude during the significant time window in panel A, showing a frontocentral distribution (N = 39). **C.** Theta activity parametrically tracked unsigned changes in feedback scores (labelled |Δ graded score|), increasing under reward and decreasing under punishment. A significant condition difference was observed within 0.2–1 s (P_FWER_ = 0.010). **D.** Topography of the effect in panel C, with condition differences localised to left frontocentral and right centroparietal electrodes. **E.** Theta amplitude as a function of the amount of upcoming changes in keystroke dynamics (ΔVⁿ), showing increased amplitude under reward and decreased amplitude under punishment between 0.2–0.9 s (P_FWER_ = 0.009). **F.** Spatial distribution of the condition difference in panel E, peaking in midline frontal and left central regions. **G.** Bayesian regression analysis. Posterior estimates of theta amplitude, averaged over the significant spatiotemporal cluster identified in C, revealed a credible interaction between PA category and reinforcement condition in relation to unsigned score changes: with increasing PA, theta increased under reward but was suppressed under punishment (posterior estimate: –2.17, 95% CrI [–4.04, –0.34]). **H.** Same as G, but for theta activity related to upcoming motor adjustments (*ΔV*^n^). Theta amplitude, averaged over the significant cluster identified in F, also showed a PA × reinforcement interaction: as PA increased, theta decreased under punishment but increased under reward (posterior estimate: –2.58, 95% CrI [–4.12, –1.07]). Dots represent posterior means, and bars denote 95% highest density intervals.

Next, Bayesian linear models revealed a credible PA × reinforcement interaction on theta modulation with unsigned changes in graded scores. As PA increased, theta amplitude in the relevant spatiotemporal cluster became more pronounced under reward but was increasingly suppressed under punishment (posterior estimate: –2.17, 95% CrI [–4.04, –0.34]; **Figure 5G)**. A similar interaction emerged for theta activity related to upcoming changes in keystroke dynamics: with increasing PA, theta was attenuated under punishment but enhanced under reward (posterior estimate: –2.58, 95% CrI [–4.12, –1.07]; **Figure 5H**).

### Dissociating the influence of reward and punishment on categorical and continuous motor decision-making

Experiment 3 examined whether the interaction between PA and reinforcement condition on learning rates and motor variability observed in Experiment 1 arose from their influence on participants’ categorical decisions—such as switching between dynamics contours (e.g., U-shape) after reinforcement—or on refining keystroke velocity within the same contour. To this end, we designed a modified reinforcement-learning task in which categorical options were explicitly displayed, reducing the action space and thus lowering task uncertainty relative to the large, continuous action space used in Experiments 1 and 2. Additionally, although skilled pianists exhibit high consistency in timing and velocity across renditions^77,78^, we examined whether individual differences in motor noise, potentially modulated by PA, contributed to the observed effects^79,80^.

A new cohort of highly trained pianists (N = 36) completed the task, which included separate baseline and reinforcement-learning phases. At baseline, participants played two new melodies with instructed constant or varying dynamics, each across 25 trials (**Figure S10**). These conditions, respectively, allowed us to evaluate unintended variability, reflecting motor noise, and total variability, encompassing both intended (exploratory) and unintended components^80^ (**Methods**).

During reinforcement learning of the same melodies as in Experiment 1 (**Figure 1**; **Figure 6A**), participants chose one of four displayed dynamics contours, including the unknown correct one, before performing the melody and refining the intensity of their dynamics within the selected contour (**Figure 6B–C**). Reinforcement feedback (reward 0–100; punishment –100 to 0) was presented in separate 100-trial blocks per condition. Pianists used this feedback to adjust both categorical and continuous aspects of performance to approach the hidden target dynamics.

**Figure 6.**
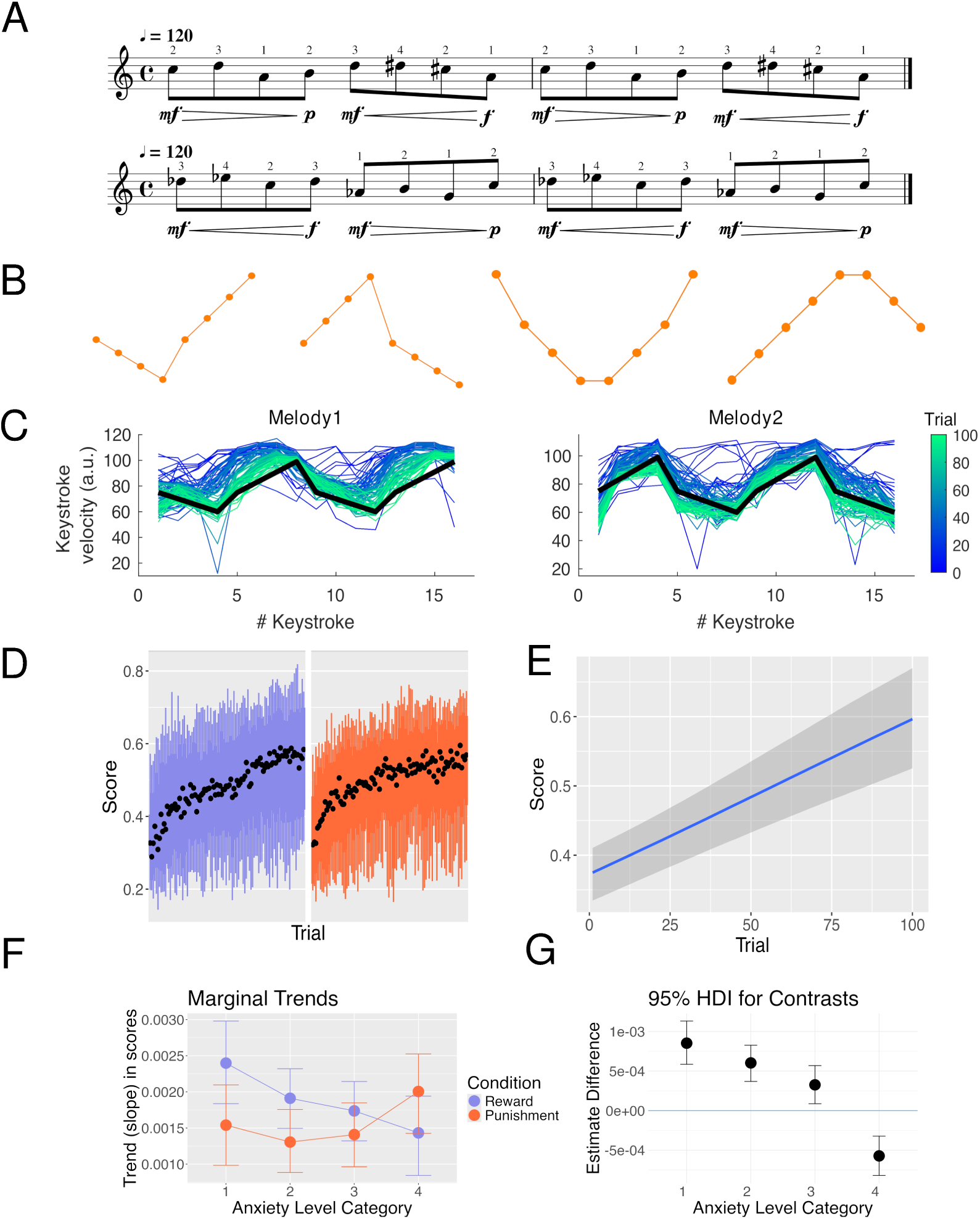
Task and Performance Analysis for Experiment 3. **A.** Right-hand melodies for the reinforcement-based performance task (same as in Experiments 1 and 2), each associated with a different hidden target pattern of keystroke velocity values, indicated in musical notation beneath the musical score. **B.** At the start of each trial, participants selected one of four displayed dynamics contours using left-hand piano keys (C2– F2), indicating both their categorical prediction of the correct contour and the one to be executed. The correct contour was 2 for Melody 1 and 1 for Melody 2. **C.** Example performance data from one participant across trials 1–100, showing keystroke velocity profiles for each melody (graded from blue to green) overlaid on the correct target dynamics (bold black line). **D.** Mean feedback scores over trials for the reward (purple) and punishment (orange) conditions, with 66% and 95% confidence intervals. The sample consisted of N = 36 skilled pianists. **E.** The posterior estimate of the overall trial effect on scores revealed a credible learning effect (slope = 0.00963, 95% CrI [0.00737, 0.01202], log-odds scale). **F–G.** Marginal trends. A credible three-way interaction between trait performance anxiety (PA) categorical level, trial, and reinforcement condition was observed. Contrary to Experiments 1–2, reward sped up learning more than punishment at low and medium PA levels. Median slope differences (reward – punishment) decreased across PA categories: low: 8.53 × 10⁻⁴ (HDI [5.85, 11.13] × 10⁻⁴); medium: 6.03 × 10⁻⁴ ([3.69, 5.36] × 10⁻⁴); medium-high: 3.26 × 10⁻⁴ ([0.873, 8.33] × 10⁻⁴). At the highest PA level, learning was faster under punishment (–5.72 × 10⁻⁴, [–8.19, –3.22] × 10⁻⁴).

We hypothesised that if reward and punishment differentially influenced categorical decisions based on PA levels, this would manifest in the rate of staying/switching between contour options. Conversely, the interaction effect could modulate the refinement of keystroke dynamics within the same contour, reflecting decision-making along a continuous scale. Learning biases might also arise from the combined effects of both categorical and continuous decision-making.

### Pianists with higher PA scores achieve more consistent keystroke velocity under instruction

At baseline, unintended variability, measured by the variance in keystroke velocity across trials ( *σ*_un_ ^2^, mean: 25.3 [SEM: 2]), was negatively associated with PA scores (Spearman *ρ* = -0.38, 95% CI: [-0.64, -0.03], *P_FDR_* = 0.025, *BF_10_* = 3.06, indicating substantial evidence for a correlation; **Figure S11**). This suggests that pianists with higher PA scores were better at maintaining consistent keystroke dynamics across trials when instructed, reflecting reduced motor noise. Conversely, intended variability 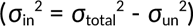 showed a nonsignificant correlation with PA (Spearman *ρ* = -0.17, 95% CrI: [-0.47, 0.14], P_FDR_ = 0.250, *BF_10_* = 0.670, anecdotal evidence for H_0_).

### PA and reinforcement condition do not modulate categorical decisions to stay or switch

During reinforcement learning, participants selected the correct dynamics contour 64.03 (0.05)% of trials. A Bayesian logistic regression revealed that higher scores (0–1 continuous range) strongly increased the probability of staying with the same contour (posterior slope = 9.15, 95% CrI [8.53, 9.78], log-odds scale). At the reference point (score = 0), the probability of staying was ∼0.06 (log-odds = –2.69, 95% CrI [–3.05, – 2.32]); at the maximum score (1), it rose to ∼0.998. There was no credible modulation of stay behaviour by reinforcement condition, PA, or their interaction (**Supplementary Materials**). Thus, categorical decisions to stay or switch between contours were driven by trial-wise feedback scores, with no credible influence of reinforcement type or PA.

#### Learning biases reflect the combined effect of categorical and continuous motor decision-making

Bayesian multilevel modelling of scores from the 64% of trials in which participants selected and played the correct contour revealed no consistent interaction between PA category and reinforcement condition on marginal trends (slopes; **Figure S12**). No credible fixed effects of PA were observed either. However, refinement of keystroke dynamics within the correct contour approached the target dynamics faster under reward than punishment (marginal trend difference: 0.00204, 95%-HDI [0.00117, 0.0029]).

Since decision-making along a continuous scale within a fixed contour category did not account for the PA × reinforcement interaction effects on learning biases in performers, the remaining analyses focused on all trials. The best-fit model (M6, LOO-CV; **Table S7; Figure S13**) showed a credible effect of trial on increasing average scores (equivalent to a gain of 19 points over 100 trials) and their precision, indicating greater consistency (**Figure 6DE**). A trial × PA × condition interaction was also observed, while PA alone did not influence slopes (**Supplementary Materials**).

Marginal effects analysis showed that reward increased learning speed more than punishment (response scale: 3.03 × 10^-4^, 95% HDI [1.34, 4.69] × 10^-4^). Further analysis of the three-way interaction revealed that, contrary to Experiments 1 and 2, learning was faster for reward than punishment at low to medium-high PA levels, with median trend differences decreasing across PA categories (**Figure 6FG**): low: 8.53 × 10^-4^ (95% HDI [5.85, 11.13] × 10^-4^); medium: 6.03 × 10^-4^ ([3.69, 5.36] × 10^-4^); medium-high: 3.26 × 10^-4^ ([0.873, 8.33] ×10^-4^). For the highest PA category, the effect reversed, showing faster learning to avoid punishment (-5.72 [-8.19, -3.22] × 10^-4^). No consistent interaction effects on marginal medians were observed **(Figure S14).**

This reversal of learning biases in Experiment 3 led us to examine whether participants adopted different learning strategies as a function of task uncertainty. We therefore conducted an exploratory post hoc analysis, adding an open question to a separate ongoing within-subject study in skilled pianists (N = 42)— originally designed for unrelated purposes—to assess whether the same high-versus low-uncertainty conditions elicited different self-reported approaches. Participants consistently indicated two distinct regimes: under high uncertainty they engaged in broad exploration, trying diverse dynamic shapes, including extremes, across the full action space, whereas in low uncertainty they relied on local exploration within a known contour, making small, targeted adjustments to refine the pattern (**Supplementary Materials**).

Collectively, the findings in Experiment 3 suggest that learning biases arose from the combined effects of categorical and continuous motor decision-making, rather than either component alone. These effects were also better explained by somatic than cognitive components of PA ( **Supplementary Materials; Figure S15**). The results remain robust even when accounting for potential modulatory effects of individual baseline levels of intended and unintended variability, neither of which exhibited credible effects on scores in this task (**Supplementary Materials**).

### Reinforcement-driven use of motor variability parallels learning biases in Experiment 3

Statistical matching analysis (**Figure 7A-D**) revealed that low scores, compared to high scores, increased motor variability over the subsequent three trials (*P_FDR_* = 0.006; Δ_dep_ = 0.71, CI = [0.62, 0.85]; **Figure 7D**). In addition, larger deviations from target velocity patterns (observing lower scores) were followed by greater reinforcement-related variability (**Figure 7E**).

**Figure 7.**
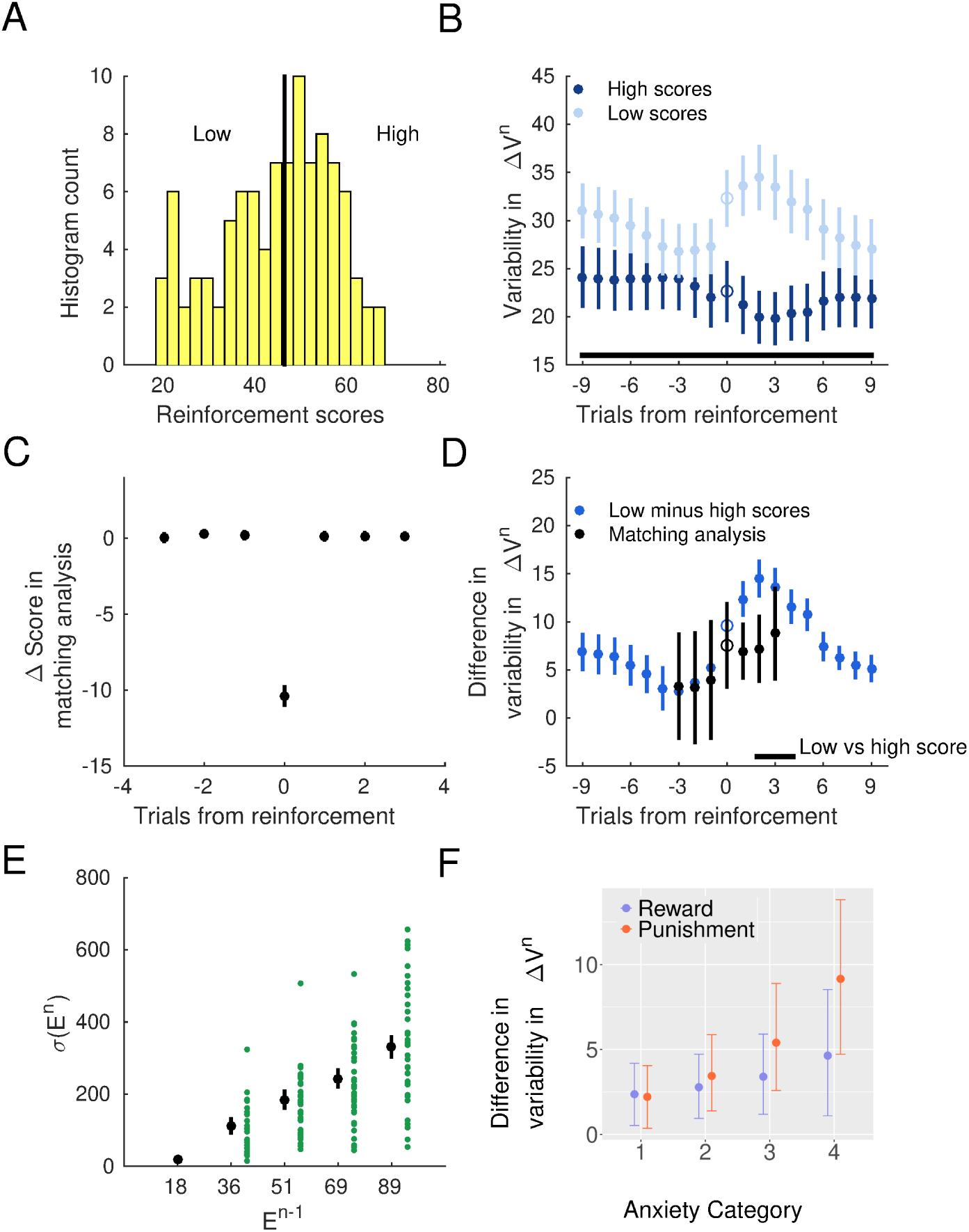
Reinforcement-related modulation of motor variability in Experiment 3. **A.** Histogram of reinforcement scores (example participant) with median split used to define low and high score conditions. **B.** Time course of motor variability (*ΔV*^n^) surrounding relatively high (dark blue) and low (light blue) score trials, aligned to the conditioned trial at position 0. Variability was greater following low scores at positions +1 to +3 (N = 36; paired permutation test; *P*_FDR_ = 0.001; significant cluster denoted by the black bar). **C.** Statistical matching analysis: trials of low and high scores (median split) were matched for surrounding reinforcement values. Black dots indicate mean performance score difference (± SEM) at each time point. **D.** Using matched trials, motor variability was significantly greater following low- than high-score trials at lags +1 to +3 (*P*_FDR_ = 0.006). Uncorrected differences from all trials (as in panel B) are shown in blue. **E.** Larger deviations from target dynamics (unsigned error *E*^n-1^) were associated with greater subsequent reinforcement-related variability, σ(*E*^n^), replicating the relationship found in Experiment 1. **F.** Bayesian linear modelling of *VarDiff* (post-low minus high score variability) showed a credible PA × reinforcement condition interaction (posterior estimate = 26.31, 95% CrI [11.34, 40.11]). Under punishment, the reinforcement-driven increase in motor variability became larger as PA levels increased, whereas under reward, posterior estimates overlapped with zero. Dots represent posterior means, bars denote 95% credible intervals.

Complementing these findings, a Bayesian Gaussian linear model of *VarDiff*—variability in keystroke velocity (*ΔV^n^*) after low minus high scores—identified a credible interaction between PA category and reinforcement condition (1.54, 95% CrI: [0.01, 3.14]; **Figure 7F**). This interaction showed that increasing PA levels were associated with a stronger modulation of variability under punishment than reward, indicating a reversal of reinforcement-dependent regulation of motor variability that paralleled the behavioural learning biases in Experiment 3.

### Reinforcement-sensitive Gaussian Process accounts for increased variability use under punishment in higher PA

Fitting the RSGP to the time series of signed errors (*e^n^*) in Experiment 3 replicated a key finding from Experiment 1: slow autocorrelations spanned ∼9 trials, whereas reinforcement effects are shorter-lived (∼3 trials). The autocorrelation kernel showed significantly greater characteristic length and output scale than the reinforcement-sensitive kernel (*l_SE_* = 9.26 [0.8] vs. *l_RS_* = 2.67 [0.3]; *P_FDR_* = 0.0002, Δ_dep_ = 0.83, CI = [0.74, 0.90]; σ_SE_ =4.20 [0.3] vs. σ_RS_ = 2.90 [0.2]; *P_FDR_* = 0.0004, Δ_dep_ = 0.70, CI = [0.58, 0.78]).

Bayesian regression showed a credible positive effect of PA on *σ_RS_* (log scale: 0.16, 95% CrI [0.03, 0.28]; Figure 8A), indicating greater reinforcement-sensitive variability with higher PA. Models including reinforcement condition or its interaction with PA performed relatively worse (**Supplementary Materials).** No effects were observed on σ_SE_.

**Figure 8.**
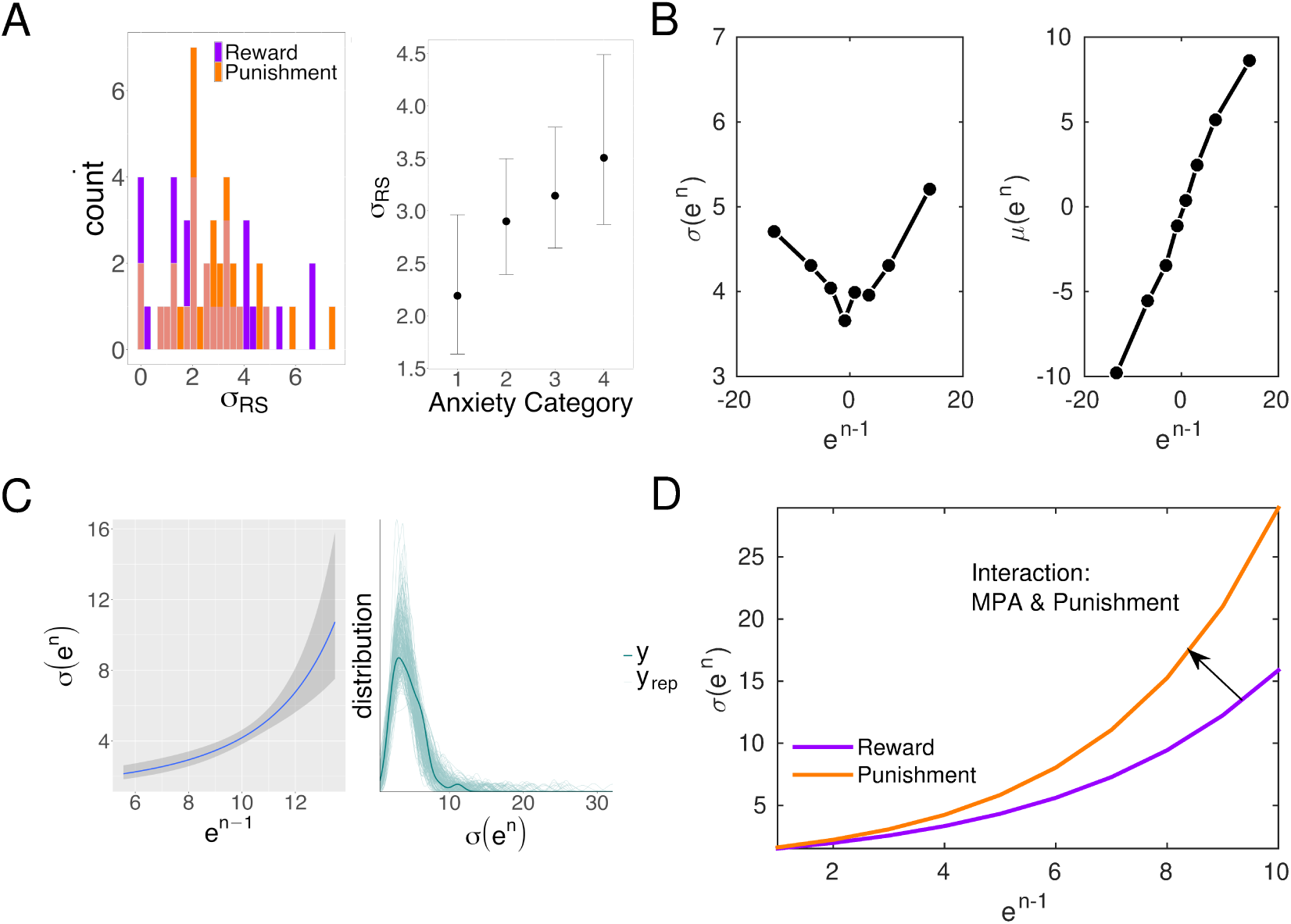
Reinforcement-sensitive Gaussian process accounts for increased motor variability under punishment in higher performance anxiety. **A.** Left: distribution of *σ*_RS_ estimates across participants by reinforcement condition (orange: punishment; purple: reward). Variable *σ*_RS_ is the output scale of the reinforcement-sensitive kernel, which captures short-term variability that is modulated by reinforcement. Right: *σ*_RS_ increased with PA category, reflecting greater contribution of reinforcement-sensitive processes to variability in individuals with higher PA levels. Estimates are in the log scale. **B.** RSGP fit to empirical data replicates simulation results: *σ(e^n^)* shows a U-shaped dependence on *e^n-1^* (left), and *μ(e^n^)* increases linearly with *e^n-1^* (right). Mean and SEM are shown; SEM values are very small and not visible. **C.** Left: exponential fit (line with 95% CrI shading) to the relationship between *σ(e^n^)* and |*e^n-1^*|. Right: posterior predictive distribution of *σ(e^n^)* values (light green lines: individual posterior draws; dark green line: empirical density). **D.** Illustration of the interaction between PA category and reinforcement condition on the exponential growth parameter *b*₂. Under punishment (orange), the growth in *σ(e^n^)* with |*e^n-1^*| was amplified in higher PA individuals relative to reward (purple), indicating increased error sensitivity and behavioural adaptation through variability. Posterior estimate: *b₂* = 0.06, 95% CrI [0.01, 0.19]

As in Experiment 1, the independent baseline noise (σ₀ = 2.23 [0.08] on average) decreased with increasing PA (log scale: -0.045, 95% CrI = [-0.119, 0.028]). This parameter correlated positively with the empirical measure of unintended variability (*σ*_un_) from the separate baseline task (*ρ* = 0.34, *p* = 0.04), supporting the partial construct validity of σ₀ as an index of individual motor noise.

Simulations using individual parameters (*l*_SE_, *l*_RS_, *σ*_SE_, *σ*_RS_, *σ*_0_) reproduced the U-shaped relationship between *e*^n-1^ and *σ*(*e*^n^), and the linear increase of *μ*(*e*^n^) with *e*^n-1^ (**Figure 8B).** Finally, the association between *total* motor variability, *σ*(*e*^n^), and the unsigned error in the previous trial, |*e*^n-1^|, was better captured by an exponential model, as in Experiment 1 (**Supplementary Materials; Figure 8C**; *b*_1_ = 1.18, 95% CrI = [0.85, 1.50]; *b*_2_ = 0.26, 95% CrI = [0.10, 0.46]). A credible interaction between PA and reinforcement condition modulated *b*_2_ (0.06, 95% CrI = [0.01, 0.19]; **Figure 8D**), increasing with PA category and under punishment relative to reward.

These findings indicate that higher PA levels enhanced the exponential growth rate of *σ(e^n^)* under punishment compared with reward, reflecting increased sensitivity to larger errors (lower scores) from the previous trial and greater behavioural adaptation through variability regulation.

## Discussion

Across three experiments, we show that predisposition to PA shapes how skilled performers learn from reward and punishment in a context-dependent manner: highly trained pianists with greater PA shifted from enhanced reward-based learning in high-uncertainty contexts—where the action space was large—to enhanced punishment-based learning when uncertainty was reduced. This reversal was paralleled by reinforcement-driven regulation of motor variability, whereby increased variability following relatively poor outcomes was modulated in the same direction as the learning biases. In Experiment 1, concurrent EEG recordings further revealed that these behavioural learning biases were mirrored by frontal theta activity parametrically encoding unsigned feedback changes and the magnitude of upcoming motor adjustments, suggesting that PA modulates prefrontal control processes engaged during reinforcement learning. Collectively, these results suggest that PA is associated with differences in how performers regulate exploratory variability to achieve performance goals, biasing the use of reward and punishment signals according to task uncertainty.

Contrary to our hypothesis, Experiments 1&2 identified positive learning biases in elevated PA levels. Decision-making studies typically report elevated punishment learning rates in mood and anxiety disorders, potentially reinforcing negative affective biases^17,18^. Consistent with this pattern, state anxiety reduces reward-based learning^24,26^ (but see ref^81^), reflecting attenuated belief updating^24,27^. By contrast, trait anxiety has been linked to faster learning and improved inference of hidden states in volatile environments^26,82^, across both reward and aversive settings. Thus, while negative learning biases have been proposed in anxiety^16–18^, the evidence shows heterogeneity across clinical and subclinical samples^17,22–24,26,82^(including null effects^83,84^), suggesting that both task demands and the populations studied can shape the expression of these biases.

The reversal of learning biases as a function of task uncertainty in our study may reflect how trait PA shapes exploration strategies in reinforcement-based motor learning. Our task was adapted from our previous reward-based motor learning research^27^, in which non-musicians used graded reward feedback to approach a hidden target solution during simple piano sequences, closely paralleling the high-uncertainty conditions in Experiments 1 and 2. That work showed that induced anxiety states reduce reward learning and attenuate motor variability. By contrast, Experiment 1 indicated that greater trait PA was associated with enhanced reward learning relative to punishment, a pattern replicated in Experiment 2. These differences between state- and trait-related effects on motor variability regulation and learning rates mirror analogous dissociations reported in categorical decision-making ^24,26^.

Trait anxiety has also been linked to enhanced directed exploration for information seeking, whereas state anxiety may diminish such exploration^85–87^, and no consistent effects have been identified for random exploration driven by decision noise. Although directed and random exploration can be dissociated only in discrete-choice paradigms that manipulate information value and decision horizon^85^—and this distinction does not readily extend to continuous action spaces—our results suggest that in large action spaces (Experiments 1 and 2), where uncovering the hidden target dynamics required broad sampling across many possible solutions, trait PA may promote reward-guided exploratory behaviour. This interpretation aligns with post hoc reports from our separate experiment, in which participants described wide-ranging exploration in the high-uncertainty condition but local, contour-guided refinement when the solution space was constrained. In Experiment 3, the reduced action space was associated with lower task uncertainty, as participants had increased information about the hidden target solution. On average, 64% of trials involved the correct categorical contour, so improvement depended primarily on using reinforcement feedback for motor refinement—stabilising movements within the chosen contour through controlled regulation of motor variability. In this more constrained setting, we observed enhanced punishment learning with higher trait PA, suggesting that high-PA individuals shift towards punishment-based learning when exploratory demands are reduced and performance hinges on refinement.

These findings also extend evidence on how anxiety dimensions influence learning. Somatic anxiety—which reflects heightened physiological reactivity—has been shown to enhance safety learning, whereas cognitive anxiety promotes threat learning^21^. In our data, somatic PA better accounted for the observed learning biases across experiments, suggesting that the physiological component of trait PA may promote reward-guided exploration under high uncertainty but shift towards punishment-based learning when the action space is constrained.

The observed learning biases were partly explained by the effects of PA and reinforcement on the regulation of motor variability. Across tasks, keystroke velocity was modulated by recent reinforcement history, increasing after low outcomes, consistent with evidence that reduced reward rates trigger behavioural adjustments^27,35,80,88^. Establishing a causal link in this association is challenging, however, because motor performance exhibits persistent correlated variation across trials^40,69,70,72,73^, which can inflate motor variability estimates^36^. To address this, we matched trials with comparable surrounding reinforcement values. In Experiment 1, higher PA was associated with greater variability regulation under reward but blunted regulation under punishment, whereas Experiment 3 showed the opposite pattern, aligning learning biases with variability regulation.

Our generative-model analysis^39^ confirmed that total motor variability—quantified as the standard deviation of the error (the deviation between produced and target dynamics)—reflected a short-lived effect of reinforcement integration over three trials, while autocorrelations persisted over 9–10 trials, consistent with prior work^39^. The latent contribution of reinforcement-sensitive variability (*σ_RS_*), likely reflecting exploratory variability, decreased with PA in Experiment 1 but increased in Experiment 3, aligning with the punishment-minus-reward differences observed in the marginal trends. Tentatively, these results suggest that punishment may exert more salient or consistent effects on the reinforcement-sensitive scaling of motor variability, as proposed in previous motor learning research^29,89–90^ (but see opposing findings in ref.^33^).

Error deviations also increased total variability on the next trial following an exponential function, and the rate of this increase was modulated by the PA × reinforcement interaction in the expected direction across experiments. These findings extend the statistical-matching results, showing that PA alters whether individuals rely more on graded reward or punishment signals when adjusting performance.

Importantly, learning biases were not explained by baseline intended or unintended variability, even though higher PA was associated with reduced motor noise (*σ*_un_). The baseline noise parameter (*σ*₀) in the generative model likewise decreased with PA and correlated with unintended variability (*σ*_un_), supporting its construct validity as an index of motor noise. Together, these findings indicate that reinforcement-dependent adaptation in skilled performance relies on contextual modulation of variability sources rather than baseline motor noise. Although optimal reinforcement learning has been proposed to require an appropriate balance between exploration variability and motor noise^34, 80,88,91^, and higher motor noise has been shown to impair learning^91^, the reduction in motor noise across PA levels in our data would, in principle, allow high-PA individuals to exploit exploratory variability more effectively. However, our results show that this advantage depends on reinforcement valence, suggesting that additional factors— potentially including working memory capacity^92,93^—may shape how PA influences reinforcement-driven exploratory variability.

On a neural level, reward and punishment learning were dissociated in the amplitude of theta-band activity. A linear convolution modelling approach^76^ revealed greater midfrontal theta 0.2–0.5 s after punishment than reward feedback, consistent with previous work^47,49^, although this effect did not relate to learning performance. More critically, a distinct theta pattern scaled parametrically with unsigned feedback changes in a reinforcement-dependent manner: during reward, theta increased with higher PA in left frontocentral and right centroparietal sites, whereas the effect was attenuated under punishment. Theta in left central and midline frontal regions also predicted the magnitude of upcoming keystroke velocity adjustments, increasing under reward and decreasing under punishment. This spatiotemporal dissociation aligned with the behavioural and computational learning biases, suggesting that PA modulates prefrontal control signals engaged during reinforcement-driven motor learning.

These findings converge with evidence that midfrontal theta encodes unsigned prediction errors and signals behavioural adjustments, consistent with its proposed role in configuring prefrontal control networks^47,49^ and with elevated frontal-midline theta in high-anxiety individuals following performance errors^51^. Theta also synchronises prefrontal and motor regions during decision-making^51^, potentially guiding action execution; in our task, this may underlie the motor adjustment effect extending to contralateral sensorimotor electrodes.

We did not model trial-wise prediction errors (PE) and cannot directly map unsigned score changes to PEs within RL or Bayesian frameworks. This may account for the absence of beta-band effects in our GLM, despite evidence linking feedback-related beta suppression to updating motor predictions^42,45,46,94^—a process dysregulated in anxiety^27^. Future work using models of hidden-state inference via PEs in continuous action spaces will be required to clarify the role of beta in PA-related learning.

To further elucidate the underlying neural circuitry, source analysis will be required to determine whether ACC and PFC activity contributes to the observed theta effects, given their roles in behavioural control^52,95^ and learning under uncertainty^14^—including via modulation of midline-frontal theta^51,52^—and their consistent functional alterations in anxiety^12,56^. Subcortical structures such as the striatum, basal ganglia, and thalamus—implicated in reward prediction-error encoding, reinforcement-based motor learning^58,60^, and the regulation of motor variability^39,59^—should also be considered to dissociate cortical and subcortical contributions to PA-related learning biases. The striatum is a particularly relevant target for future work, as it carries both valence-specific and valence-nonspecific prediction-error signals, with dorsal regions responding selectively to unexpected reward and ventral regions (along with PFC) responding to both unexpected reward and unexpected punishment^96^. A key direction for future work is to characterise how interactions between PFC, ACC, and motor circuits support reinforcement-driven motor adaptation in skilled performers, and how these interactions may be altered in high-stakes contexts that trigger PA.

While the inclusion of multiple experiments in expert performers strengthens the robustness and validity of our findings, the focus on pianists limits generalisability to other expert populations. Future studies should examine whether similar effects emerge in other high-performance domains where PA is prevalent. Additionally, while we assessed learning biases as a function of trait predisposition to PA, it will be important to examine how these biases manifest under experimentally induced PA, particularly across different forms of uncertainty, and to incorporate measures of intolerance of uncertainty, a hallmark of anxiety^24^ to clarify their contribution to learning biases in performers.

In sum, our findings demonstrate that predisposition to PA in skilled individuals modulates learning from reward and punishment in a context-dependent manner. As uncertainty increases, faster learning shifts from punishment to reward, driven by the active regulation of motor variability and modulation of theta-band activity. Exploratory evidence suggests that high-uncertainty contexts are perceived as more aversive, particularly among individuals with greater intolerance of uncertainty, offering a potential affective mechanism underlying this shift. These findings offer important insights into how trait PA shapes learning biases and identify neurocomputational processes through which high-stakes, uncertain environments may impair expert performance.

## Materials and Methods

### Experiment 1

#### Demographics

Forty-two participants were recruited for this study, aiming for a sample size of 40 to achieve the desired power level (simulation-based Bayesian power analysis: **Supplementary Materials)**. One participant was excluded for not adhering to the task procedure. The final sample (N = 41, 23 females, 18 males; age range: 18–66, M = 29.2, SEM = 2; 32 self-reported right-handed, 8 left-handed) comprised pianists with at least six years of formal piano training, with proficiency in sight-reading sheet music, advanced musical technique, and an understanding of music dynamics. On average participants had 19.16 (SEM 2.0) years of training and performance, and were currently playing an average of 11.95 (SEM 1.9) hours per week.

Participants did not have a history of neurological or psychiatric conditions and were not currently taking medication for anxiety or depression. Due to faulty EEG recording in one participant, the EEG analysis sample consisted of 40 participants (23 females, 17 males).

All participants gave written informed consent, and the study protocol was approved by the local ethics committee at the Department of Psychology, Goldsmiths, University of London. Participants were compensated with £35, with the possibility of increasing this sum up to £45 depending on their task performance.

The Kenny music performance anxiety (K-MPAI) inventory evaluates cognitive, behavioural, and physiological components commonly associated with MPA and other anxiety disorders^97^, demonstrating high consistency across cultures and various musician populations^98^. It consists of 40 items rated on a 7-point Likert scale (0 “strongly disagree” to 6 “strongly agree”). Scores range from 0 to 240, with values above 160 considered above average. Previous factor analysis of the K-MPAI in professional musicians identified sub-scores associated with proximal somatic anxiety and negative cognitions, which we used to assess the dissociation between somatic and cognitive dimensions of PA on learning in our study^99^. We also administered the trait subscale of the Spielberger State-Trait Anxiety Inventory (STAI Form Y-2^100^; Spielberger, 1983), assessing more generalised anxiety (See **Supplementary Materials)**. Participants completed the questionnaires at the beginning and end of the session, respectively.

#### Procedure

Upon arrival, participants were seated at a digital piano (Yamaha P-255) and familiarised themselves with two short right-hand melodies (Melodies 1 and 2; **Figure 1A**). They memorised the melodies using the indicated fingering and practised them with a metronome (120 bpm) for 3–10 min to ensure consistent tempo. The metronome was removed during the main task. After participants could play both melodies from memory (five consecutive error-free renditions), EEG equipment was fitted (30–45 min).

The main task involved discovering the hidden target dynamics of the two melodies—specific keystroke-velocity patterns differing from natural phrasing. We chose melody dynamics as the target variable because the rendition of dynamics can vary widely among performers, introducing a degree of ambiguity in how they are executed during musical performances. Participants were informed that the target dynamics would deviate from conventional expressive patterns and were encouraged to explore different dynamics guided by feedback. They were shown example dynamic contours (e.g., crescendo, diminuendo, mixed shapes) illustrating potential solutions (**Figure S1**).

Each participant completed 100 trials per reinforcement condition, presented in separate reward (0–100 scores) and punishment (–100 to 0) blocks (**Figure 1B**). Feedback reflected the proximity between the performed and target keystroke-velocity vectors, computed using an exponential decay function applied to the square root of the summed absolute velocity differences between the participant’s MIDI keystrokes and the target dynamics, adjusted to the predetermined scale in each reinforcement condition. MIDI velocity values ranged 0–127, and the digital piano volume was fixed at a medium level across sessions. In the reward condition, scores indicated monetary gains up to £5; in the punishment condition, scores represented losses from £5, using identical scoring functions. To ensure consistency, the researcher followed a scripted protocol when delivering instructions.

Trials with pitch errors received the lowest possible score (0/-100) and were excluded from analysis (6.98 [0.74]% reward; 7.24 [0.85]% punishment; *P* = 0.77, *BF*_10_ = 0.18). Trained pianists are highly proficient at detecting self-made pitch errors^101,102^, supporting that lowest feedback scores from such trials were not misattributed to incorrect dynamics. Each trial began with a cue, followed by a 7 s window to perform the melody, and a 2 s feedback display. The task was implemented in Visual Basic with MIDI and parallel-port interfaces. Condition order and melody–reinforcement mapping were pseudorandomised and counterbalanced across participants.

#### Stimulus materials

Melody 1 and Melody 2 (**Figure 1A**) were composed specifically for this study. These two short melodies, in a 4/4 time signature, consist of a pattern of 8 quavers repeated twice over 2 bars.

The 8-note dynamic pattern was repeated to form a musically coherent 16-note phrase suitable for pianists, and participants were informed of this. The melodies were designed with the following criteria in mind: (i) they were to be played with the right hand only; (ii) their performance would not present technical challenges and would require minimal shifts in hand or finger positioning to reduce movement artifacts affecting EEG recordings; (iii) the melodies were to be atonal; (iv) they would be presented to the participants with no dynamics indicated (**Figure 1A**); (v) the melody’s hidden target dynamics would not be a trained musician’s initial guess (**Figure S1**), necessitating exploration of dynamics to infer the solution (**Figure 1C**). The target dynamics (**Figure 1A,C**) disrupted the traditional beat structure of the 4/4 time signature, whereby the first and third beat are strong, and the second and fourth are weak. Crescendos, decrescendos and accents that were included as part of the melodies’ target dynamics furthermore clashed with the natural flow and direction of the melody.

#### EEG, ECG, and MIDI Recording

EEG and ECG signals were recorded using a 64-channel EEG system (ActiveTwo, BioSemi Inc.), following the extended international 10–20 system, placed in an electromagnetically shielded room. During the recording, the data were high-pass filtered at 0.1 Hz. Vertical and horizontal eye movements (EOG) were monitored by electrodes placed above and below the right eye and at the outer canthi of both eyes, respectively. Additional external electrodes were placed on both left and right mastoids to serve as initial references upon importing the data into the analysis software (data were subsequently re-referenced to a common average reference, see below). The ECG was recorded using two external channels with a bipolar ECG lead II configuration: the negative electrode was placed on the chest below the right collarbone, and the positive electrode was placed on the left leg above the hip bone. The sampling frequency was 512 Hz.

As in our previous EEG studies with trained pianists^101,103^, participants were instructed to minimise upper body and head movements during trial performance and outcome processing, focusing movement solely on their fingers. This was facilitated by our stimulus material, which was designed to avoid shifts in hand or finger positioning, thereby reducing movement artifacts. Participants were informed that they could briefly move between trials if needed, and that they could initiate the next trial at their own pace.

Performance was recorded as MIDI files using the software Visual Basic and a standard MIDI sequencer program on a PC with Windows XP software (compatible with Visual Basic and the MIDI sequencer libraries we used). To run the behavioural paradigm and record the MIDI data, we used a modified version of the custom-written code in Visual Basic that was employed in similar paradigms in our previous studies^27,103^. This program was also used to send synchronisation signals in the form of transistor–transistor logic (TTL) pulses—corresponding with onsets of visual stimuli, key presses, and feedback scores—to the EEG/ECG acquisition PC.

#### Bayesian analysis workflow of performance data

We modelled trialwise performance scores using Bayesian beta regression models (brms package, v2.21.0^104,105^; R, v4.3.2). Beta regression is appropriate for bounded outcomes in (0,1), parametrised by the mean (*μ*) and precision (*ϕ*). Reward scores (0–100) were rescaled to 0–1 by division; punishment scores were shifted by +100 and then divided by 100. The empirical score distribution avoided boundary values, satisfying beta-model assumptions. Error trials (fixed score 0 after rescaling) were excluded, as described in the motor variability analyses.

To build and evaluate these models, we followed the principled Bayesian workflow^61,106,107^, proceeding through prior specification, prior predictive checks, model fitting, and posterior predictive evaluation. Our initial model (M1) captured fixed effects of reinforcement condition, trial, and their interaction. Priors were set using domain knowledge from prior reward/punishment learning studies^17,21^: a positive trial effect on *μ*, a positive condition × trial interaction (faster punishment learning), and no baseline difference between conditions (***Supplementary Materials***). The *μ*-intercept prior (0.2 on the log-odds scale; ≈0.55 on the score scale via *plogis*) used a *σ* = 0.1; remaining *μ*-coefficients used Gaussian priors centred on their expected value (smaller *σ* = 0.01). For *ϕ*, Gaussian priors were centred at 0 (*σ* = 0.01), except for the intercept (centred at 2.5, *σ* = 1) and trial (small positive mean) to reflect expected growing score consistency across trials. Prior predictive simulations (sample_prior=“only”) confirmed that M1 generated plausible ranges for minimum, maximum, and mean scores (**Figure S4**) and values were consistent with our domain expertise^106,107^.

We then expanded to increasingly complex models (M2–M6; **Table S1**), adding random intercepts (M2), and random slopes (M3). The most complex model included a three-way condition × trial × PA interaction with subject-level intercepts and slopes (M6). Following recommendations by Bürkner and Charpentier (2020)^108^ on using ordinal regressors in Bayesian regression models, to assess PA effects , we used a monotonic predictor for ordered PA categories (quartiles: 70, 107, 125; range 18–169). The STAI-Trait ranged 20–65 (median 40), approximating population norms (reference median norm is 35, and values above 45 are considered very high^109^). See further details in ***Supplementary Materials***.

Random-effect correlations in M3 and M6 used an LKJ(2) prior (class = “cor”). The remaining parameters, related to monotonic effects of PA, were set to default priors. We checked the prior predictive distribution in each model, similarly to M1, which confirmed their adequacy^110^.

Models were fitted with 4 chains, 5,000 iterations each (1,000 warm-up; 16,000 post-warm-up samples). Model comparison was performed employing the leave-one-out cross-validation of the posterior log-likelihood (LOO-CV) with Pareto-smoothed importance sampling^65^. We selected the model with highest expected log pointwise predictive density (ELPD), and verified that the absolute mean difference in ELPD (*elpd_diff*) between the two best-fitting models was at least 4 and larger than twice the standard error of the difference (2*se_diff*). Where *elpd_diff* < 2×*se_diff*, the more parsimonious model was chosen. Convergence was verified using R-hat < 1.01 and effective sample sizes (ESS) exceeding recommended thresholds^111,112^. Posterior predictive checks confirmed adequate fit^61^.

For the winning model, we report population-level posterior medians and 95% credible intervals (CrI) for relevant parameters (full population-level estimates are provided in ***Supplementary Materials).*** *C*redible differences were defined by CrIs excluding zero. To quantify learning-rate differences as a function of PA, reinforcement, and trial, we used *emtrends* (emmeans v1.10.1), which provides marginal trends for the slope coefficient in non-linear models; trends were transformed to the response scale using *regrid=“response”*. Marginal medians were obtained with *emmeans*. The same modelling approach and priors were applied in Experiment 2.

#### Analysis of Motor Variability

Motor variability was quantified from the 16-dimensional keystroke-velocity vector (8 × 2 notes) using a scalar metric of trial-to-trial change (see e.g. Banca et al 2023^68^):

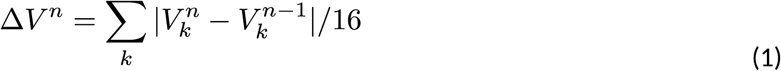

Here, *ΔV^n^* denotes the normalised sum of absolute differences in keystroke velocity between consecutive trials (*n-1* and *n*) across all *k* positions in the melody. *ΔV^n^* thus captures the scalar magnitude of change in the multidimensional velocity vector from one trial to the next (**Figure 1C**). Variability was estimated as the variance of *ΔV^n^*values across running windows of five trials, following Dhawale et al. (2019)^36^.

To determine whether participants increased variability following poor outcomes relative to good ones^35,36,68^, we analysed motor variability separately for low and high scores (median split per participant and melody; **Figure 3A**). Because time series of behavioural performance, including errors in timing, press angles, and endpoint reaching, exhibit medium- to long-range (persistent) autocorrelations^40,69–73^, which confound the assessment of causal relationships between variability and performance^36,39,73^, we applied a validated statistical matching procedure^36^ to control for these effects. This method matches low- and high-scoring (“conditioned”) trials preceded and followed by similar “matched” reinforcement values (**Figure 3C**), isolating reinforcement-driven changes in exploratory behaviour from autocorrelation effects, thereby providing a more accurate estimate of the causal influence of reinforcement on subsequent motor variability.

The matching analysis was applied separately for each melody. We then averaged the task-relevant variability in *ΔV^n^* following conditioned trials across melodies, contrasting it between low and high score conditions using paired permutation tests (5,000 permutations). The results demonstrated significantly higher motor variability following low scores compared to high scores at positions +1 to +3 after the conditioned trial (see ***Results***, **Figure 3D**).

Importantly, our analysis was carefully designed to prevent incorrect trials from contaminating or confounding the results (**Supplementary Materials**).

To investigate whether learning biases towards reward or punishment were influenced by variations in the causal relationship between reinforcement and motor variability across PA levels, we specifically employed a Bayesian Gaussian linear model. This model included fixed effects of PA category, reinforcement condition, and their interaction to analyse changes in task-relevant variability (*VarDiff*). Specifically, we examined *VarDiff* values from the first three positions following conditioned trials, as these positions exhibited significant variability differences in our statistical matching analysis (**Figure 3D**). See further details in **Supplementary Materials**.

#### Generative model of behaviour: reinforcement-sensitive Gaussian Process

We employed a reinforcement-sensitive Gaussian Process (RSGP) model to characterise trial-wise variability in keystroke velocity errors as a function of reinforcement history. This generative model extends the reward-sensitive Gaussian Process framework developed by Wang et al. (2020)^39^ for analysing variability in reinforcement-based motor tasks. Gaussian Processes are probabilistic models that infer continuous functions from noisy observations by defining a prior distribution over functions, with dependencies between samples (here trials) captured by a covariance function^113,114^. The RSGP uses a composite kernel, combining a standard squared exponential kernel (K_SE_) to model slow autocorrelations in behaviour with a reinforcement-sensitive kernel (K_RS_), which modulates trial-wise covariance as a function of reinforcement scores. Specifically, *K*_RS_ is a squared exponential kernel with zero covariance for unrewarded (here low outcome) trials. This formulation ensures that only rewarded (high outcome) trials contribute to *K*_RS_, tightening behavioural coupling to recent successes and enabling increases in observed motor variability following unsuccessful trials. The RSGP also includes a time-independent baseline noise term with an identity matrix scaled by *σ^2^*(**Figure 4A; Supplementary Materials** ).

The RSGP was fitted to trial-wise error values (*e^n^*), defined as the expectation of the difference between the produced (***V*^n^**) and target (***T***) keystroke velocity vectors, averaged across the 16 keystroke positions for each melody rendition:

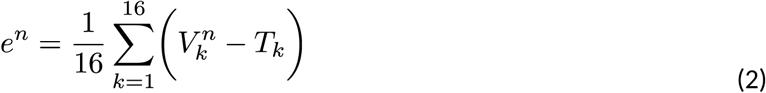

We use lowercase *e^n^* to distinguish this signed error metric from the error metric illustrated in **Figure 3E**, which is based on the norm of vector differences and is always positive. The model estimated the variance and mean of *e^n^* on each trial based on prior values and reinforcement history. The kernels were parametrised by characteristic length scales (*l_SE_*and *l_RS_*) and output scales 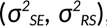, along with baseline noise *σ^2^*, all of which were inferred via Bayesian inference^114^.

Following Wang et al. (2020)^39^, we validated the model through simulations, testing its ability to recover known hyperparameters (**Supplementary Materials**). When fitting the model to empirical data, we estimated *l_SE_, l_RS_, σ_SE_, σ_RS_*, *σ*_0_ separately in each participant and condition to capture individual differences. We used Matlab code for RSGP simulation and model fitting from ref.^39^ (https://github.com/wangjing0/RSGP).

To examine the effects of PA and reinforcement condition on model parameters, we implemented Bayesian regression models with fixed effects. Model selection was based on leave-one-out cross-validation (LOO-CV), and parameter credibility was assessed using posterior predictive checks.

See further details in **Supplementary Materials**.

#### EEG preprocessing and analysis

EEG preprocessing was done in MATLAB R2020b using the toolboxes EEGLAB^115^ and FieldTrip^116^. EEGLAB was used to import the files and filter the data, applying a 50 Hz notch filter to remove the power line noise. Data were downsampled to 256 Hz.

For the Independent Component Analysis (ICA), we applied a high-pass filter at 1 Hz to improve ICA decomposition^117^. The data were then segmented into epochs from -2 to 2 seconds locked to the outcome trigger, thereby minimising the presence of any potential movement artifacts that could emerge between trials. During trial performance and outcome processing, participants had been instructed to minimise movements. ICA was run in FieldTrip, using the runICA algorithm^118^, which combines the Infomax algorithm^119^ with the natural gradient learning rule^120^. After IC decomposition, we applied the resulting ICA weights to the 0.1 Hz-filtered version of the epoched data, as recommended by the EEGLAB developers. Artifacts related to eye blinks, eye movements (saccades), and cardiac artifacts, if present, were removed (3.7 on average, range 2-5).

Manual inspection of the epochs was then performed to remove any remaining artifactual epochs, such as those affected by muscle artifacts (reflected in high-frequency fluctuations^121^). This resulted in a total of 78.6 (SEM 3) and 71.3 (SEM 3) clean epochs left for the analysis of the punishment and reward conditions, respectively. In cases where a channel was faulty throughout the epoch inspection, interpolation was used to replace this channel with the average signals from neighbouring channels. This happened with 1-3 channels from 5 participants. One EEG dataset was excluded due to large muscle artifacts during the feedback presentation interval, leaving N = 39 datasets for analysis.

To model neural EEG responses to feedback scores and motor variability, we used linear convolution models for oscillatory responses^76^. This approach extends the classical general linear model (GLM) from fMRI analysis to time-frequency (TF) data and has been widely applied in EEG and MEG research ^122,123^. It enables trial-by-trial assessment of TF response modulation by a specific explanatory regressor while controlling for the effects of other included regressors.

In all convolution analyses, each discrete and parametric regressor was convolved with a 20th-order Fourier basis set (40 basis functions: 20 sines and 20 cosines). This configuration enabled the GLM to resolve TF response modulations up to ∼8.7 Hz (20 cycles/2.3 s; ∼115 ms). The discrete regressor was modelled by convolving this chosen basis set of functions with delta functions encoding the timing of the feedback events, commonly referred to as stimulus input.

We compared three GLMs including regressors for feedback onset, upcoming keystroke adjustments (scalar variable *ΔV^n^*), and trial-wise scores, which were modelled as (1) graded scores, (2) unsigned or (3) signed score changes from the previous trial. Collinearity analyses between regressors in each GLM design ( **Supplementary Materials**) showed strong correlations between graded scores or signed score changes and *ΔV^n^*, but minimal overlap for unsigned score changes. Accordingly, the selected convolution GLM included the unsigned score and *ΔV^n^* as parametric regressors, alongside the discrete regressor for feedback onset.

The selected GLM was applied to concatenated epochs spanning -0.5 to 1.5 s around the feedback event, using Morlet wavelets for time-frequency (TF) analysis in 4–30Hz, thus covering the theta, alpha and beta ranges. We conducted this analysis using SPM12 software (http://www.fil.ion.ucl.ac.uk/spm/), adapting original code by ref.^122^, as used in^26,124^.

Statistical analysis of sensor-level time-frequency images used cluster-based permutation testing in the FieldTrip Toolbox^116,125^ (1000 permutations). We averaged TF activity across frequency bins within each band (theta, alpha, beta). Temporal intervals of interest for statistical analyses were selected based on previous research^27,47,48,124^: 0.2–1.5 s for parametric regressors, 0.1–0.6 s for the feedback onset regressor. We controlled the family-wise error rate (FWER) at 0.05 (two-sided tests, effects considered if *P_FWER_* < 0.025).

#### Statistical analysis

Complementing the Bayesian multilevel and non-nested models in our study, when assessing within-subject differences in a variable (e.g., variability estimates between trials of low and high scores), we implemented paired sign permutation tests with 5,000 permutations. In those cases, we additionally provided a non-parametric effect size estimator, the probability of superiority for dependent samples (Δ_dep_), which is the proportion of all paired comparisons in which the values for condition B are larger than for condition A^126^. 95% confidence intervals (CI) for Δ_dep_ were estimated with bootstrap methods^127^. To control for multiple comparisons arising from, for example, different permutation tests conducted on neighbouring trials or across several interrelated variables, we implemented the adaptive false discovery rate control at level *q* = 0.05.

### Experiment 2. Replication study

#### Demographics

A sample of 18 pianists (15 females, 3 males; 17 self-reported right-handed; age range: 18– 28, mean age = 21.1, SEM = 0.8) completed the same experimental task as in Experiment 1. Participants undertook the task, which was programmed in Python, at the Sony Computer Science Laboratory (Tokyo), using a KAWAI VPC1 digital piano with keystroke velocity in range 0–127. The same inclusion and exclusion criteria pertaining to Experiment 1 were applied for this experiment.

Written informed consent was obtained from all participants, and the study protocol was approved by the local ethics committee at Sony Corporate, Tokyo. Participants received a monetary remuneration for their participation. They received a fixed amount of 3000 JPY, which could increase by an additional sum of 4000 JPY (2000 JPY for reward, 2000 JPY for punishment conditions) depending on their task performance.

As in Experiment 1, to assess trait aspects of PA, we used the Japanese version of the Kenny MPA Inventory, and the trait subscale of the STAI-Y2.

#### Bayesian Data Analysis

Analysis of the evolution of scores over time as a function of PA and reinforcement condition was performed exactly as in Experiment 1. The PA scores in this sample were split into four partitions through the quartile boundary values 114, 131, and 144 (range 84–180; T-STAI scores ranged 39– 55).

### Experiment 3

This experiment employed a modified version of the paradigm used in Experiments 1 and 2, designed to isolate categorical decision-making from decisions made on a continuous scale in skilled pianists.

#### Participants

Thirty-six pianists (N= 36, 31 females, 5 males; age range: 19-54, M= 25.83, SD=1.3; all self-reported right-handed) were recruited for this experiment. Sample size estimates were adjusted for this experiment based on the findings from Experiments 1&2 (**Supplementary Materials**). They had not completed Experiment 2 and were naive to the task setting (Sony CSL, Tokyo). The inclusion and exclusion criteria were the same as in Experiments 1 and 2. All participants provided written informed consent, and the study protocol received approval from the local ethics committee at Sony CSL, Tokyo. Participants were compensated with 3000 JPY, with the possibility of increasing this sum up to 4000 JPY depending on their task performance. Consistent with Experiments 1 and 2, PA levels were assessed using the Japanese version of the K-MPAI questionnaire, and the trait subscale of the STAI-Y2 was also administered. Participants completed the questionnaires at the start and end of the session, respectively.

#### Paradigm and procedure

The paradigm comprised a baseline variability assessment phase and a reinforcement learning phase.

Baseline phase: Participants performed two simple right-hand melodies (Melodies 3 and 4; 4/4 time, eight quavers repeated twice; **Figure S10**) across 25 trials each to assess unintended and intended variability of keystroke velocity. After a 5-minute familiarisation, pianists played Melody 3 with consistent dynamics to measure unintended variability and Melody 4 with deliberately varied dynamics to measure intended variability. The order of conditions was pseudorandomised and counterbalanced across participants.

Reinforcement learning phase: Participants learned the hidden dynamics of the same two melodies from Experiments 1–2 through reward (0–100) or punishment (–100 to 0) feedback, but now with an added categorical action-selection stage (**Figure 6A–B**). Each trial began with a 3-s contour-selection screen displaying four dynamic patterns (chosen via piano keys C2–F2), followed by melody performance (∼8 s) using the selected contour. Participants refined their dynamics based on feedback score presented for 2 s, aiming to maximise gains (reward) or minimise loses (punishment) in each condition by approaching the hidden target solution.

The target dynamics differed from Experiments 1–2: each melody’s hidden pattern corresponded to one of the displayed contours or its inverted counterpart (patterns 1–2 for Melodies 1–2, respectively; **Figure 6A– B**) and deviated from natural phrasing. Thus, crucially, while participants could infer the correct contour after several trials, they still needed to use reinforcement (reward or punishment) to refine their performance and maximise scores by approaching the target solution.

#### Analysis of baseline variability

Variability in the task-relevant dimension, keystroke velocity, was assessed using the variance of the 25-trial distribution of keystroke velocity values. This index was first calculated for each of the 16 keystrokes of the melody and then averaged across all positions. We separately measured unintended variability (*σ*_un_ ^2^), representing motor noise, and intended (exploratory) variability, obtained from the total and unintended variability measures: 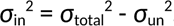.

#### Bayesian Performance analysis in Experiment 3

Following the analysis of categorical decisions using a Bayesian logistic regression, we used Bayesian multilevel Beta regression models to assess learning across the 64% of trials where participants chose the correct categorical dynamics contour. In addition, we conducted this analysis using the total dataset, including trials where participants chose an alternative contour. In both cases, the models were built as in Experiments 1 and 2. We retained the original priors from Experiment 1 rather than updating them based on the posterior estimates. This decision was driven by our task modifications in Experiment 3, which could render the previous posterior estimates less applicable. The initial priors, validated by prior predictive checks **(Figure S4**), provided a suitable starting point under the altered experimental conditions. The quartile values of PA scores in this sample were 114 (Q1), 131 (median), and 144 (Q3), with a range of 85 to 182. The trait STAI values ranged from 32 to 62.

#### Analysis of motor variability and RSGP modelling

In Experiment 3, the analysis of reinforcement-driven modulation of motor variability was conducted similarly as for Experiment 1. In addition, the same RSGP modelling approach and analysis as described for Experiment 1 was implemented.

### Data availability

Behavioural and EEG data are publicly available at the Open Science Framework (OSF) repository, https://osf.io/w7y5k/. Analysis code to reproduce the main analyses is publicly available at OSF, https://osf.io/w7y5k/.

## Acknowledgements

The research was partially funded by the Strategic Research Fund of Goldsmiths University of London (MHR), by the Economic and Social Research Council (ESRC) and the South East Network for Social Sciences (SeNSS) through grant ESRC grant number ES/Y001834/1 (AEH, MHR), and by JST CREST (JPMJCR20D4) and JST Moonshot R&D (JPMJMS2012) (SF, TO, YH). The authors thank Jenny Halseid for support in part of the data collection (Experiment 1, Goldsmiths University of London).

